# Target-based *de novo* design of cyclic peptide binders

**DOI:** 10.1101/2025.01.18.633746

**Authors:** Fanhao Wang, Tiantian Zhang, Jintao Zhu, Xiaoling Zhang, Changsheng Zhang, Luhua Lai

## Abstract

Cyclic peptides have become a new focus in drug discovery due to their ability to bind challenging targets, including “undruggable” protein-protein interactions, with low toxicity. Despite their potential, general methods for *de novo* design of cyclic peptide ligands based on target protein structures remain limited. Here, we developed CYC_BUILDER, a reinforcement learning based fragment growing method for efficient assembly of peptide fragments and cyclization to generate diverse cyclic peptide binders for target proteins. CYC_BUILDER employs a Monte Carlo Tree Search (MCTS) framework to integrate seed fragment exploration, fragment fusion based peptide growth, structure optimization, evaluation and peptide cyclization. It supports peptide cyclization through both head-to-tail amide bond and disulfide bond formation. We first validated CYC_BUILDER on known protein-cyclic peptide complexes, demonstrating its ability to accurately re-generate cyclic peptide binders in terms of both sequences and binding poses. We then applied it to design cyclic peptide inhibitors for TNF*α*, a key mediator in inflammation-related diseases. Among the nine experimentally tested designed peptides, four showed potent binding to TNF*α* and inhibited its cellular activity. CYC_BUILDER provides an efficient tool for cyclic peptide drug design, offering significant potential for addressing challenging therapeutical targets.

## 1 Introduction

Many diseases have been found to involve multifactorial aspects and complex networks of protein-protein interactions (PPIs).^1–3^ These interfaces are usually large with buried solvent accessible surface areas ranging from 1500 to 3000 Å^2,4,5^ which are difficult to be covered by small molecules (usually with solvent accessible surface areas between 150-500 Å2).^4,6^ In contrast, peptides are well-suited to block PPIs.^7^ Peptides offer several advantages over traditional small-molecule drugs. Their biosynthesis and solid-phase synthesis methods are well-established,^8^ and their degradation products, amino acids,are minimally toxic. Despite these advantages, linear peptides face significant challenges, including poor membrane permeability,^9,10^ rapid degradation and reduced binding strength due to entropic losses upon binding. Cyclization of linear peptides has emerged as a highly effective solution to overcome these limitations.^4,11–14^ A number of cyclic peptide drugs have been already used to treat diseases,^15^ such as T-cell lymphoma, lupus pneumonitis, and excessive blood loss.^16–18^

Machine learning has significantly advanced small molecule drug and protein design.^19,20^ Early efforts have extended to the design of cyclic peptides. The availability of cyclic peptide conformation databases, such as CREMP,^21^ has enabled efficient cyclic peptide conformation sampling using deep learning algorithms, such as StrEAM, ^22,23^ RINGER,^24^ have achieved efficient sampling of cyclic peptide conformations. Several groups have modified Alphafold to optimize the backbone structure of free cyclic peptides or cyclic peptide-protein complex and design sequences for cyclic peptide binders.^25–28^ However, due to the limited number of crystal structures of cyclic peptide-protein complexes, *de novo* design of cyclic peptide ligands solely based on the target protein structures remains largely unexplored.^29^

This study introduces a target protein structure based cyclic peptide binders design method, CYC_BUILDER, using Monte Carlo Tree Search (MCTS) framework and reinforcement learning algorithms.^30^ Cyclic peptides are grown in the protein binding site using fragment-based building blocks to give stabe and natrual-like conformations. By incorporating reward functions that account for both binding strength and structure rationality, CYC_BUILDER efficiently generates cyclic peptide binders for diverse protein targets and exhibits good generalization capability. Compared to the anchor extension strategy proposed by Hosseinzadeh et al using Rosetta^31–38^ and other deep learning methods such as AfCycDesign^25,26^ and RFpeptide,^28^ our algorithm explores high-quality cyclic peptide scaffolds rapidly and guides the intermediate structures towards multiple cyclization patterns (both head-to-tail and disulfide bonds) using potential score functions. Sequence and backbone diversity of the generated cyclic peptide ligands are also promising. We have applied this method to design and experimentally validated cyclic peptide binders for tumor necrosis factor *α* (TNF*α*), a key player in inflammation related diseases, and obtained potent peptide inhibitors.

## 2 Overview of CYC_BUILDER

CYC_BUILDER offers a general fragment growing strategy for structure based *de novo* design of cyclic peptide binders. Based on the fragment database extracted from the proteinprotein interface, our program efficiently constructs head-to-tail or disulfide cyclized peptides (See Section 5.3) targeting protein-protein interfaces, including both surface and pocket regions. Monte Carlo Tree Search drives peptide growth and cyclization, progressively refining fragment selection for optimal conformations without reliance on complex structural training data. With advanced scoring methods, the tool assesses conformation stability, binding affinity, and cyclization propensity, utilizing grid-based interaction scoring and enhanced intramolecular hydrogen bond analysis. CYC_BUILDER excels in creating peptide binders with strong affinity and diversity, making it well-suited for complex protein-protein interactions.

The main framework consists of 4 modules, including fragment sampling, fragment growth, structure optimization and backbone closure. The growing process starts from a seed fragment, which serves as the root node of the Monte Carlo tree. The seed peptide fragment can be extracted according to the hotspots of an existing protein or peptide binder, or it can be generated by the Seed seeking algorithm (See Section 5.2). Originating from the root node, each growth action in our model corresponds to the sampling of a new set of fragments from the tripeptide structure library and the fragment fusion to the current peptide. To enhance the sampling efficiency, we categorized the tripeptide fragments in the library based on bockbone conformation, the orientation of middle residue, and the polarity and size of residues (20 natural amino acid residues were classified into 7 categories, see Section 5.1 and Figure 7). The probability matrix for sampling fragments is initialized according to the type of the fragments. The sampled fragments are then assembled onto the already constructed peptide using a flexible assembly algorithm.

Each growth action (conducted by fragment splice algorithm) that creates a new leaf is evaluated by the scoring function, which contains four terms: the binding energy, the properties of the interaction interface, the rationality and stability of the cyclic peptide structure, and the propensity for cyclization. Subsequently, sample probability matrix and state values are updated through backpropagation. In the state of the current leaf node, the model assesses whether the termination conditions can be meet. If the peptide meets the termination criteria, the model executes a cyclization operation. Otherwise, the peptide continues to grow. (See Figure 1 and Section 5.5).

**Figure 1:**
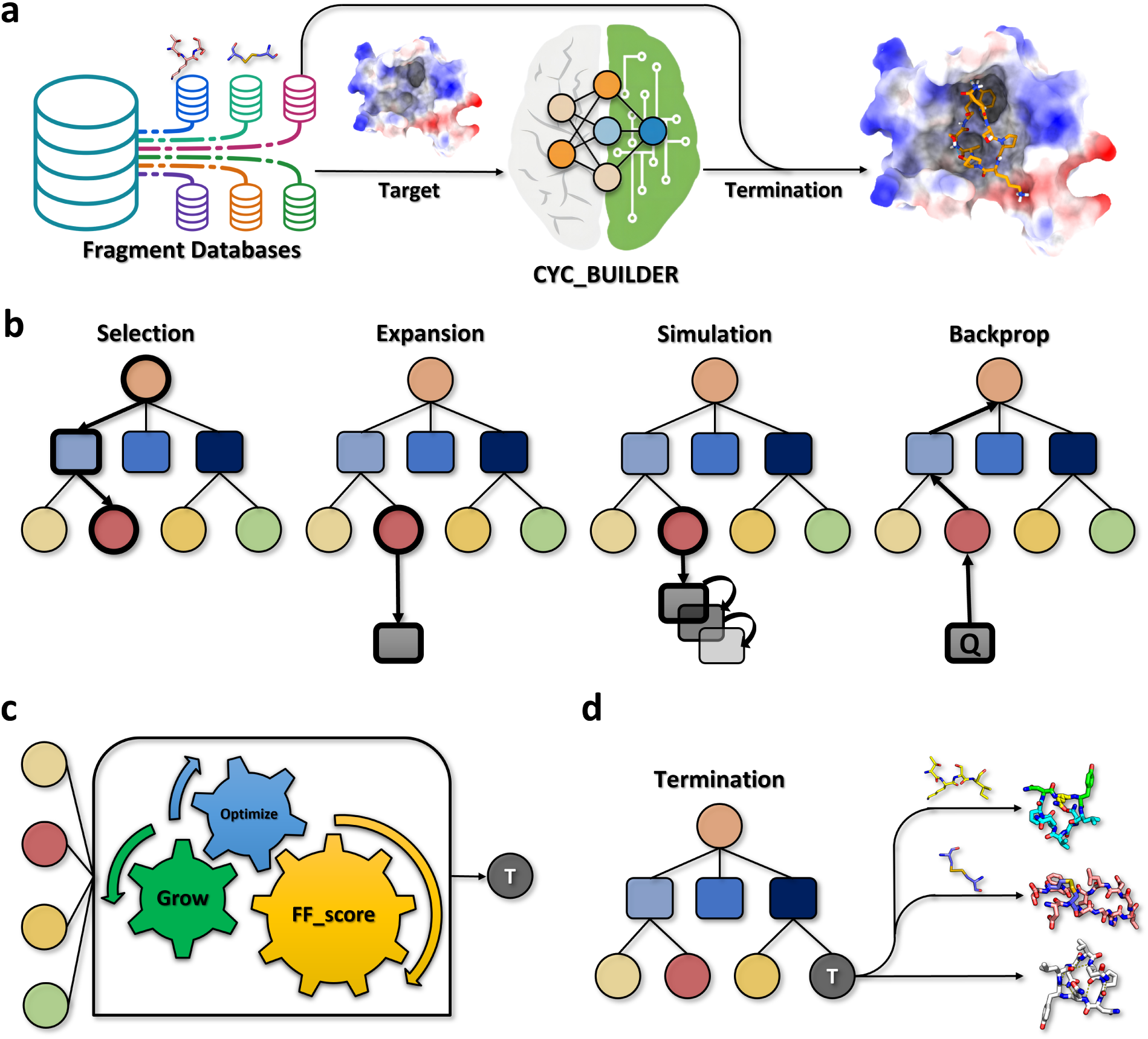
Overall framework of the CYC_BUILDER. The algorithm will first generate an anchor fragment (which can also be predefined). Then it samples from the fragment database and combines with the anchor fragment to perform the growth algorithm. It interacts with the MCTS framework to perform backpropagation, updating the sampling probability matrix and policy, and cyclizing the polypeptide until it meets the cyclization criteria to end the search. FGL represents oligopeptide fragment library, CP represents cyclic peptides, and *π* represents sampling probability matrix.

In the traditional MCTS algorithm, the rollout/playout operation in the simulation process is carried out by the Monte Carlo method. That is, it extends the leaf node from the current state until the search termination condition is met, referred to as an episode.^39,40^ Then, the score of the simulation result is backpropagated to the current node, stored as a part of the reward of this simulation in the current State. The sampling scores of the current state in combination with information such as the number of visits were used to evaluate the value of this action (denoted by Q). To ensure that the simulation component of the model converges rapidly to reasonable values, CYC_BUILDER growth module adopts a time-dependent (TD) algorithm^41–43^ rollout method. The simulation explores the next state for the benifit of avoiding the issue of episodes failing to terminate (peptide cyclized). Moreover, the sampling matrix and scoring weights are optimized by policy gradients along the growing process. Max sampling depth and a maximum child nodes number can be specified by users (See Section 5.4 for details).

In the structure optimization module, during the growing process, the peptide backbone was perturbed first and then the side chains were repacked. Efficient scoring functions were developed for the evaluation of peptide-protein target affinity, backbone conformation stablity and cyclization tendency (See Section 5.5). The output complex structure of generated cyclic peptides are optimized and filtered using Rosetta.^31^

## 3 Results

### 3.1 Regeneration test

We trimmed the ADCP cyclic peptide redock test-set^44^ which contain the X-ray crystal structures of proteins complexed with cyclic peptides for testing whether our program CYC_BUILDER can generate cyclic peptide binders with native interactions and similar binding affinity. Cyclic peptides containing 6-20 natural amino acid residues in the redock test-set were collected and redundant structures with peptide sequence similarity ≤ 40% (calculated by Biopython^45^) were removed. For complexes containing disulfide-cyclized peptides, we only collected disulfide-cyclized peptides with one disulfide bond and the residues between the two cystein residues are at least 50% of the total number of residues. The trimmed dataset CYCB-19C contains 19 entries, including 7 head-to-tail and 12 disulfide cyclic peptides, which was used for the regeneration test (See Table S3 for details). For each case, seed fragments were obtained from the seed generation algorithm (See Section 5.2, binding motif is defined as the residues within 10 Å of the geometric center of the original ligand) and inputted into the generation algorithm. The cyclization method (head-to-tail or disulfide) is restricted to the same type as the corresponding peptide in the dataset. CYC_BUILDER executed a single generation run for each target protein in the data set with a max sampling depth of 5 and a maximum child nodes number of 1000. To verify the stability of the generated cyclic peptide ligand binding and evaluate binding affinity, we conducted redocking tests using ADCP on the CYCB-19C dataset. In the ADCP redocking process, 2.5 million docking poses are sampled and 50 parallel searches were performed for each case. The results are shown in Table 1, for different types of test complexes.

**Table 1:** Affinity, L-RMSD and F*_NC_* by ADCP redock assays of designed cyclic peptide (average value of top1 cluster in a single round growth) ligand compared with original ligand. SS and CN stand for peptides cyclized with side-chain disulfide bonds and head-to-tail backbone amide bonds, respectively.

| PDB_ID | Designed |  |  | Original |  |
| --- | --- | --- | --- | --- | --- |
| | Affinity (kcal/mol) | L-RMSD (Å) | $F_{NC}$ (%) | Affinity (kcal/mol) | L-RMSD (Å) |
| 1sfi (CN) | −22.4* | 1.7* | 89.8 | −12.9 | 1.9 |
| 3av9 (CN) | −11.9* | 0.8* | 72.1 | −6.8 | 1.1 |
| 3p8f (CN) | −20.2* | 1.2* | 98.9 | −13.6 | 1.5 |
| 3zgc (CN) | −11.6* | 0.9* | 96.5 | −10.7 | 0.9* |
| 4k1e (CN) | −22.1* | 2.0 | 84.1 | −12.8 | 1.7* |
| 4kel (CN) | −14.7* | 3.5 | 75.3 | −12.4 | 2.5* |
| 5xn3 (CN) | −11.3 | 2.7* | 32.7 | −15.1* | 3.7 |
| 1smf (SS) | −22.3* | 1.0* | 66.7 | −21.9 | 1.3 |
| 1vwd (SS) | −17.7* | 3.3 | 95.0 | −16.7 | 2.0* |
| 2ck0 (SS) | −18.1 | 1.9* | 77.9 | −21.5* | 2.1 |
| 3g5v (SS) | −19.8 | 0.8* | 54.1 | −25.4* | 0.9 |
| 3p72 (SS) | −25.2* | 0.9* | 80.3 | −25.1 | 1.6 |
| 3wnf (SS) | −21.2* | 1.7 | 72.2 | −18.5 | 1.1* |
| 4ib5 (SS) | −25.6 | 2.5 | 86.7 | −27.1* | 1.6* |
| 4m1d (SS) | −23.4* | 2.6 | 65.4 | −18.7 | 1.8* |
| 5djc (SS) | −16.7 | 3.1 | 78.6 | −26.7* | 1.1* |
| 5eoc (SS) | −17.1 | 2.5 | 52.9 | −22.5* | 1.1* |
| 5th2 (SS) | −24.7 | 2.0* | 82.6 | −29.9* | 2.1 |
| 5vb9 (SS) | −17.2 | 2.5 | 53.3 | −27.3* | 1.3* |

<sup>a</sup> \* means better results;

The Top1 redock binding affinity and L-RMSD (ligand C*α* RMSD with receptor aligned) are shown in Table 1. ADCP performs well in docking natural cyclic peptides, with RMSD values typically below 2 Å, achieving a success rate of 78.9%. In 63.2% and 52% of the cases, the designed cyclic peptide ligands outperform the native ones in redock affinity and L-RMSD, respectively. Distributions of the total L-RMSD and affinity data is shown in Figure 2a, 2b and Figure 3a,3b. As shown in Figure 2c,2d, the native ligand of 1sfi, labeled as 1sfi gt, is displayed at the top of Figure 2d. The generated peptide was able to recall the key hydrogen bond interaction and hydrophobic interaction at position K1. Additionally, compared to the native ligand, s0d19 cn formed more hydrophobic interactions with the surrounding pocket. Among the SS test systems, 41.6% of the SS test systems generated peptides with binding energies better than the natural ligands as well as 41.6% performed better in L-RMSD. The disulfide-containing structures generated by the model cover a larger potion of the target site interface, forming additional hydrophobic and electrostatic interactions. For instance, as is shown in Figure 3c and 3d, in the original ligand, L6 directly contacts the negatively charged interface, while in the designed structure, an R4 residue is introduced to interact with this region, which provides a more rational interaction. Additional examples of regeneration are provided for reference in Figure S6. At the same time, we also calculated the recall rate of native interactions, F*_NC_* (Fraction of Native Contacts), for the generated structures. In the redocking tests, the Fnc was consistently maintained above 50%.

**Figure 2:**
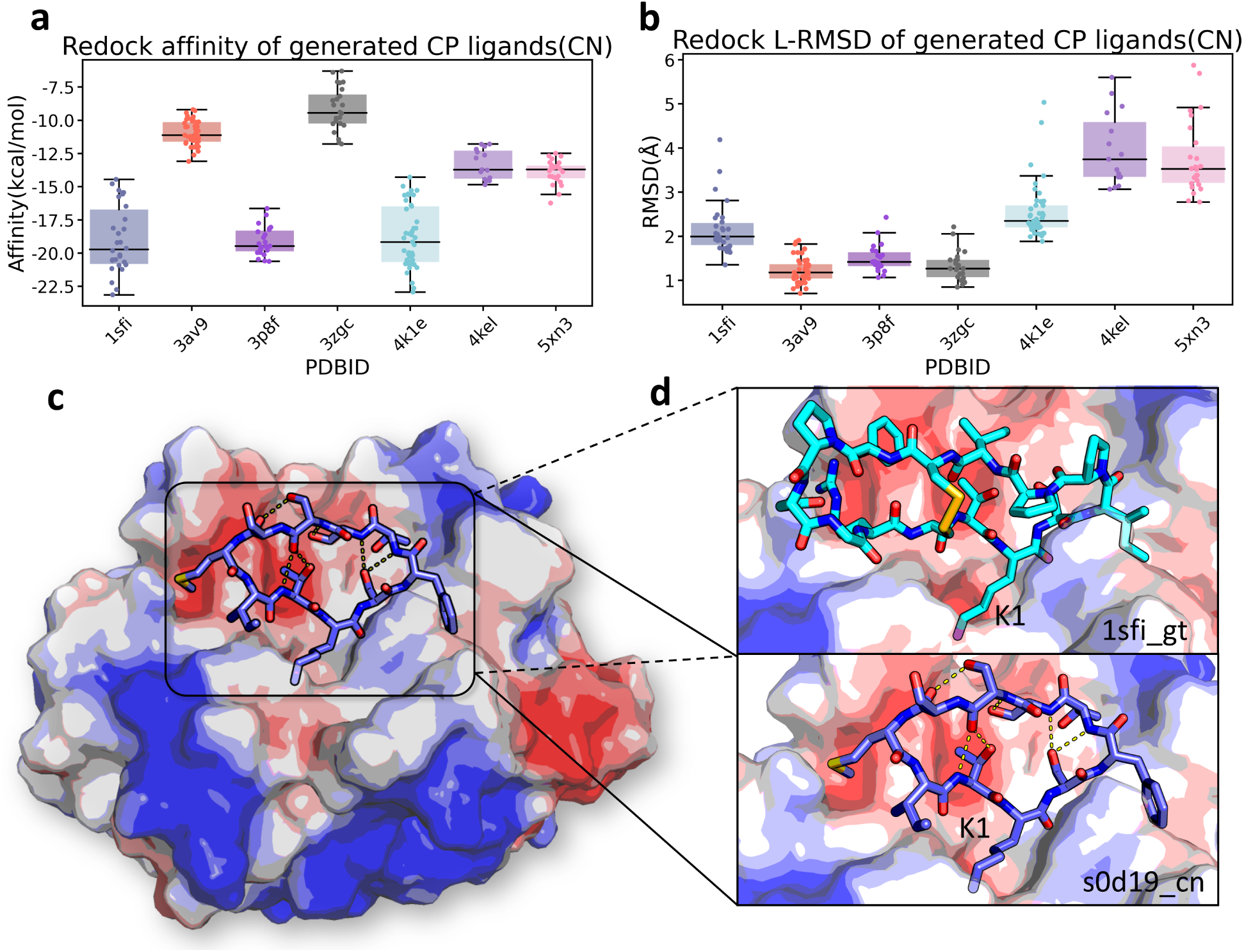
Regeneration result of the CN cases. a. Box plot of affinity of ligands generated in CN (Native ligand cyclizes through head-to-tail backbone amide bonds) tests. b. Box plot of L-RMSD of ligands generated in CN tests. c. Complex structure of s0d19 cn with Bovine *β*-trypsin (PDBID:1sfi). d. Zoom-in structure of 1sfi gt and s0d19 cn

**Figure 3:**
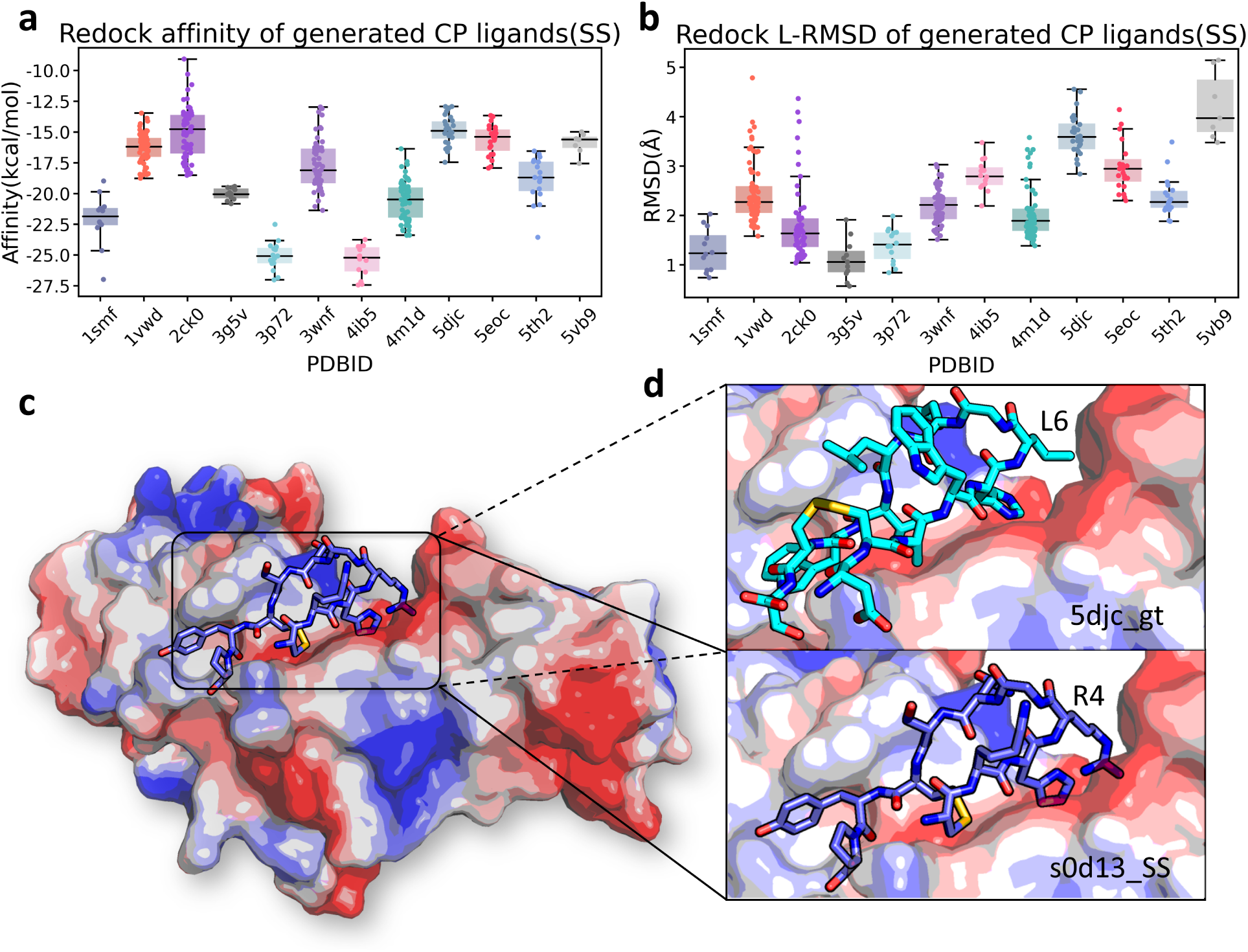
Regeneration result of the SS cases. a. Box plot of affinity of ligands generated in SS (Native ligand cyclizes through side-chain disulfide bonds) tests. b. Box plot of L-RMSD of ligands generated in SS tests. c. Complex structure of s0d13 ss with Ig gamma-1 chain C region (PDBID:5djc). d. Zoom-in structure of 5djc gt and s0d13 ss.

These results demonstrate that our model not only produces good generation outcomes but also exhibits strong generalization capability across different target proteins.

### 3.2 *De novo* generation of cyclic peptides targeting TNF*α*

Tumor necrosis factor alpha (TNF*α*) is a cell signaling protein (cytokine) involved in systemic inflammation and is one of the cytokines that make up the acute phase reaction.^46,47^ We have selected human-TNF*α* complexed with a nanobody VHH2 (PDB id: 5m2j)^48^ inter-face as the input of the seed-seeking module. We performed an alanine scan on the interface of TNF*α* and VHH2, calculating the ddG of the interacting residues. Through a sliding window approach, we selected every three consecutive oligopeptide fragments with ddG ≤ −2.0 kcal/mol to add to the seed library (See Figure S7a). This was followed by generation through CYC_BUILDER’s seed-seeking algorithm, selecting the top 10 seeds with the lowest ddG (evaluated by the Rosetta interface analyzer^49^) and best *Score_seed_* to add to the seed library. We executed the cyclic peptide growth program starting from each of the seeds in the library, with a maximum sampling depth set to 5 and a maximum child node number of 500, and generated a total of 100,000 head-to-tail cyclic peptide decoys, with peptide length between 7 and 15.

We analyzed the quality of generated cyclic peptide binders for TNF*α* using Rosetta. As shown in Figure 4, 38% structures exhibited an interface area greater than 1500 Å2, while 26% had individual ddG values below -48 kcal/mol. We selected the results with interface area greater than 1500 Å2, ddG less than -40 kcal/mol, fewer than five unsatisfied hydrogen bonds on the interface, and peptide total energy less than 0 kcal/mol for MD simulation filtering process.

**Figure 4:**
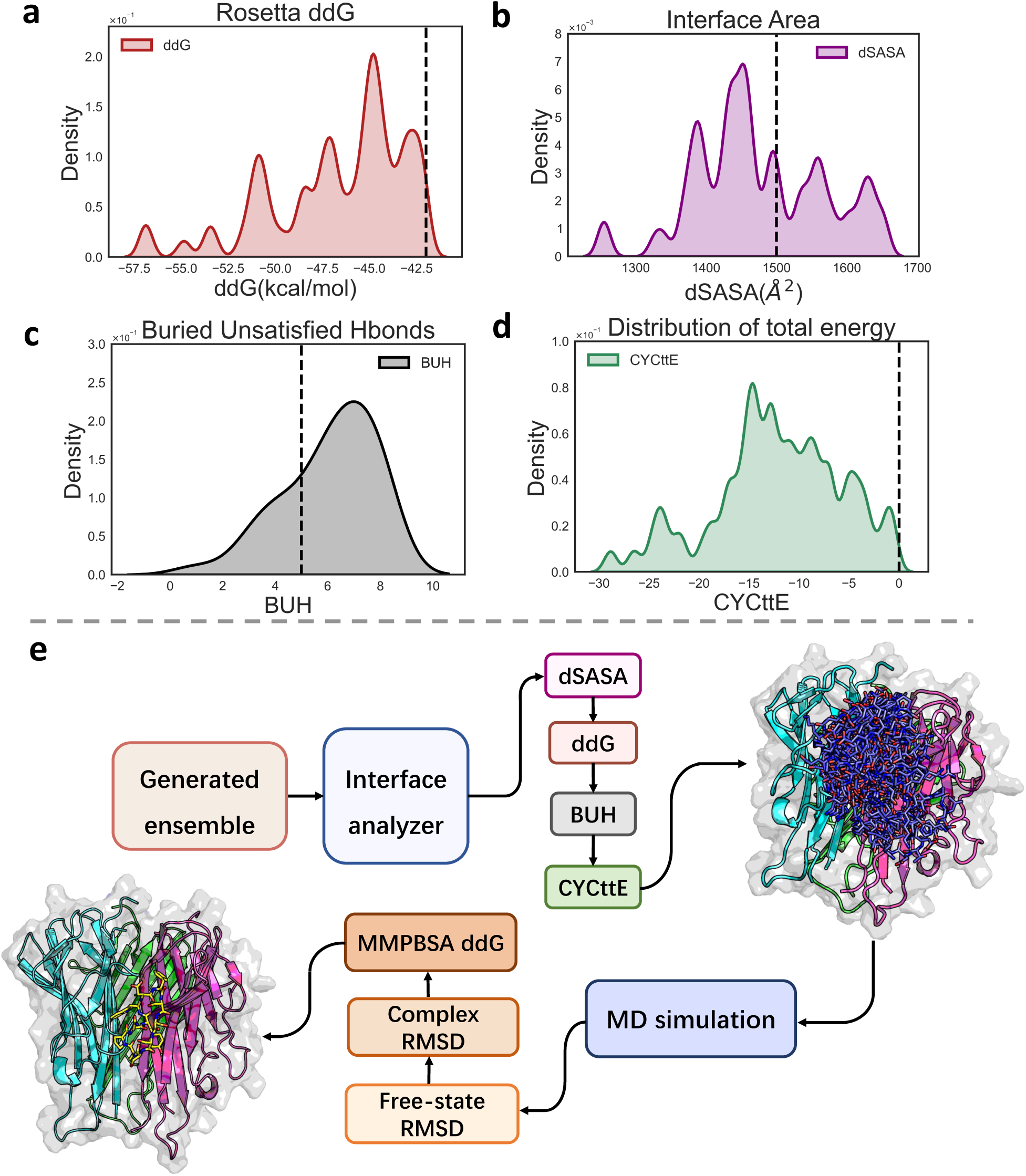
Rosetta interface analyzer metrics and free state peptide MD trajectories. a. Rosetta binding ddG of ligands generated, b. buried interface area, c. the number of buried unsatisfied Hbonds for generated ligands, d. total energy distribution, e. Flow chart of filtering processes.

We performed 100 ns molecular dynamic (MD) simulations for each of the selected cyclic peptides. We calculated the average C*α* RMSD between the simulation conformations in the last 30ns and the designed structure after the backbone superposition to evaluate the conformation change of the designed free cyclic peptide during the MD simulation. The results show that 42.5% of the cyclic peptides had a backbone RMSD less than 1.8 Å (Figure S8). We selected 37 peptides with small conformation change (RMSD ≤ 1.5 Å) for the next complex structure simulation. These designed target-peptide complex structures were subjected to 100ns MD simulations. We evaluated the average binding conformational changes in the last 30ns by calculating the backbone L-RMSD and the RMSD of the interface residues (iRMSD) between the simulation conformation and the designed structure after the alignment of the target protein structure. Residues with buried surface area in the designed structure greater than 30 Å^2^ or the burial ratio greater than 30% were assigned as interface residues. The designs with large binding conformation change in the MD simulation (iRMSD ≤ 2.5 Å) were not considered for the experimental test. Subsequently, we calculated the binding free energy of the complex structure using the MMPBSA method with the last 30ns MD trajectories. The designs with calculated binding free energy MMPBSA ddG (See ddG column in Table 2) ≤ -35 kcal/mol or the per-residue binding free energy MMPBSA ddG normalized by sequence length ≤ -3 kcal/mol (See ddG/n column in Table 2) were selected (FHT01-06) for synthesis and binding experiment test in the first round.

**Table 2:**
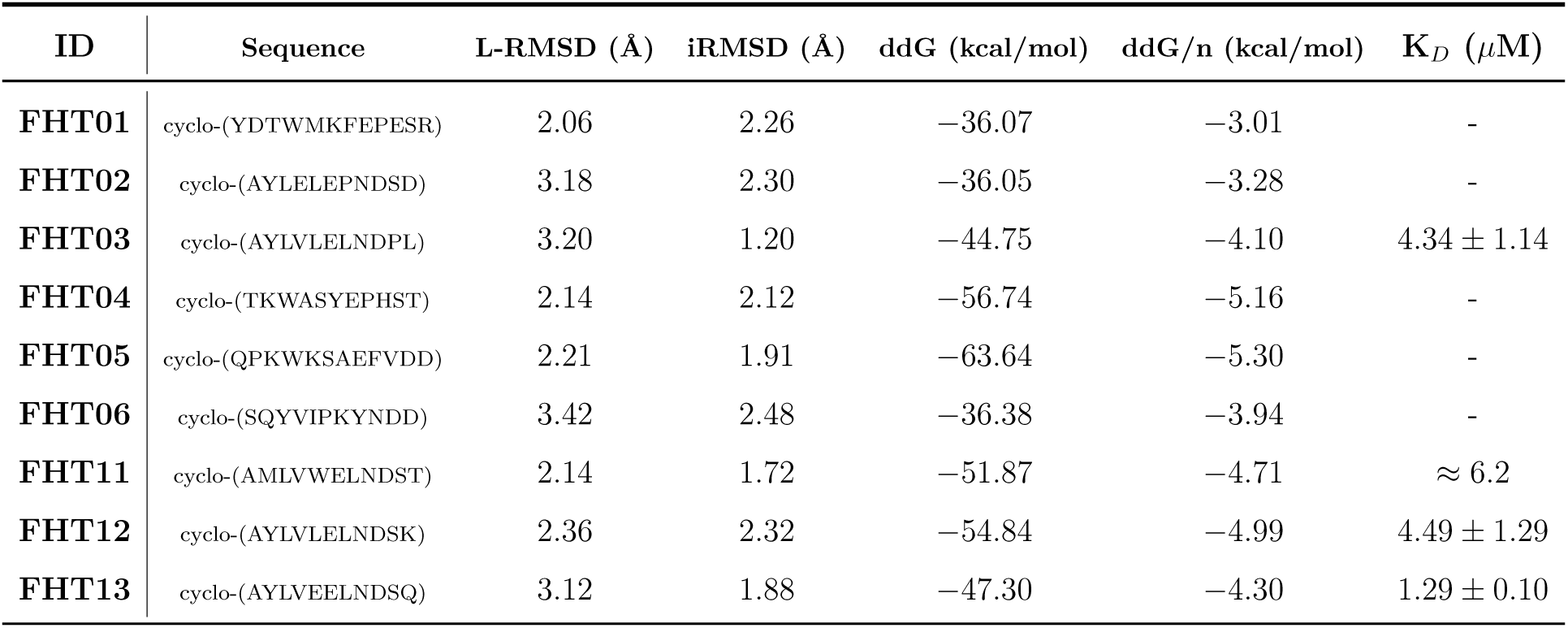
Sequence and binding properties of experimentally tested cyclic peptides binders designed for TNF*α*.

We first tested the binding strength of the designed peptides to TNF*α* using Surface Plasmon Resonance (SPR).The purified TNF*α* protein was immobilized on a CM5 chip and the cyclic peptides were used as the analytes. One of the six cyclic peptides, FHT03, showed significant binding to TNF*α*, with a K*_D_* of 4.34 *µ*M fitted by a 1:1 Langmuir binding model (Figure 5a). In luciferase-based cell activity assays, different concentrations of FHT03 were incubated with TNF*α* and significant inhibition of TNF*α*-TNFR interactions was observed with increasing concentrations (Figure 5b). Then real-time qPCR quantified the time-dependent expression of three TNF*α*-induced early-responses genes (A20, IL6 and CXCL2) and one late response gene (NK4) were performed. As a control, cells were stimulated using 4 ng/ml TNF*α*. Under this condition, the majority of cells were able to be activated to initiate the NF*κ*B signaling pathway. For inhibition test, we added 25 *µ*M FHT03 and incubated it with 4 ng/ml TNF*α* for 12 hours and measured downstream gene expression levels of the cells collected at different stimulation time points (1, 2, 3, 6, 10h, See Figure S11).^50^ For both early and late genes, FHT03 induced smaller peak expression compared to TNF*α* alone, further confirming the inhibitory effect of FHT03 on TNF*α*-induced NF*κ*B signaling pathway. In other words, our designed FHT03 not only showed strong binding to TNF*α*, but also exhibited inhibitory effects on TNF*α* at the cellular level. The energy decomposition results from MMPBSA showed that the key residues contributing to the binding with TNF*α* are R31 and R32 from chain B, as well as Y87 from chain C, with contributions to the binding energy of -20.92 kcal/mol, -9.94 kcal/mol, and -4.68 kcal/mol, respectively (Table S4). To further confirm the interaction of FHT03 with TNF*α*, three TNF*α* mutants (R31G, R32V, and Y87S) were prepared (Figure 5e), which were assumed to disrupt key hydrogen bonds, salt bridges, or hydrophobic interactions between TNF*α* and FHT03. The CD spectra of the mutants are similar to the wild-type proteins indicating that their secondary structure was not significantly altered (Figure 5c). SPR binding test showed that the FHT03 cyclic peptide did not interact with three TNF*α* mutants (Figure 5d), validating the binding position of FHT03 cyclic peptide.

**Figure 5:**
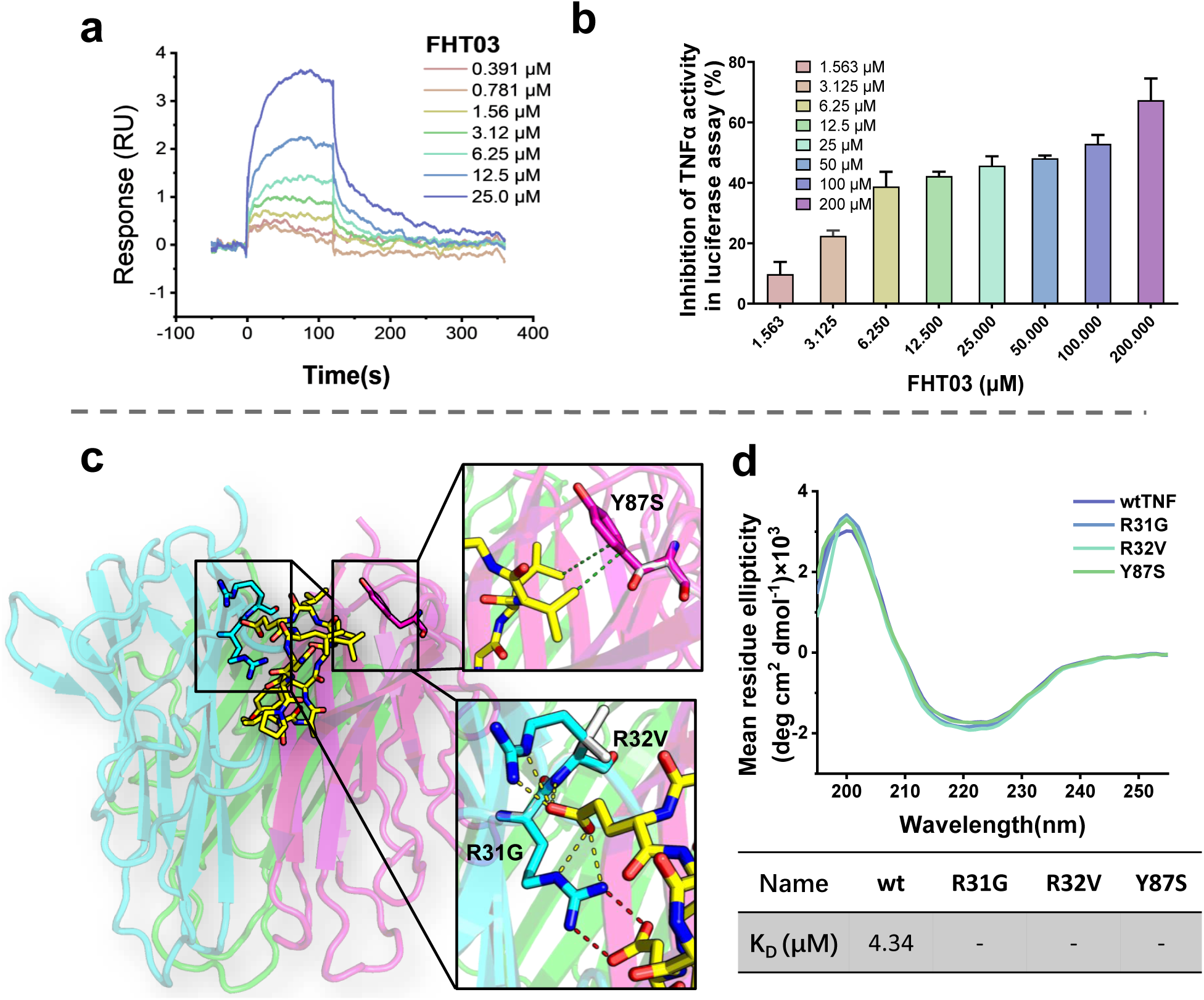
FHT03 experimental results and binding site analysis. a. SPR binding curves between TNF*α* and FHT03. b.Bar chart of cell activity from luciferase experiments corresponding to the four cyclic peptide ligands, c. schematic diagram of the binding site of FHT03 with TNF*α*, and on the right is a schematic diagram of the interactions between the three mutants and FHT03, d CD spectra of mutants R31G, R32V and Y87S, table of SPR binding between TNF*α* mutants and FHT03, “-” means non-binding.

Subsequently, since FHT03 exhibits preliminary activity and correct binding site, we relaxed the RMSD filter threshold for free-state cyclic peptide MD simulations to 1.8 Å, further screening the resulting cyclic peptides to identify those generated from the same seed fragment, Seed*_fht_*_03_ (as shown in Figure S7b), and applied the same complex structure MD screening process. This filtering process resulted in the identification of three cyclic peptide ligand structures: FHT11, FHT12, and FHT13. Notably, all three new cyclic peptides showed good binding activity with TNF*α* in SPR experiments. The K*_D_* values for FHT11, FHT12, and FHT13 are 6.2*µ*M, 4.49*µ*M and 1.29*µ*M, respectively (Figure 6a). In luciferase experiments, FHT12 and FHT13 were able to induce similar inhibitory effects at lower concentrations compared to FHT03 (Figure 6b). FHT12 and FHT13 were able to significantly reduce the expression of all the four NF*κ*B target genes induced by TNF*α* stimulation. (Figure 6c and 6d). It suggests that the binding of FHT12 and FHT13 to TNF*α* may represent a successful case of a cyclic peptide designed to inhibit TNF*α* activity. The cyclic peptide ligand generation model we developed was successfully applied to the design of TNF*α* inhibitors and showed remarkable response in SPR binding assay and significant inhibition of TNF*α* activity in cell activity assay. The strong binding of our designed cyclic peptide to TNF*α* confirms the reliability of our designed cyclic peptide ligand.

**Figure 6:**
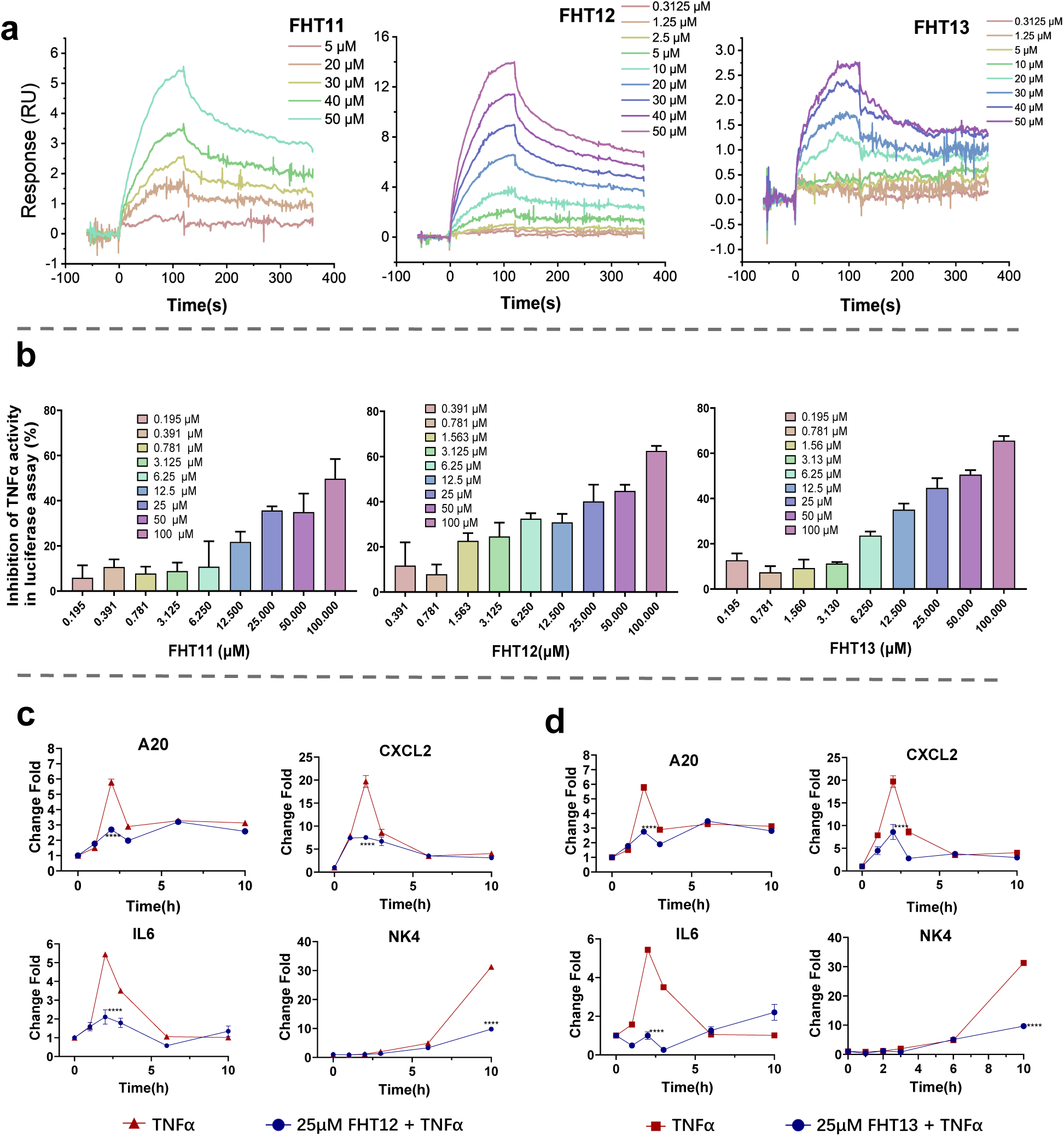
Experimental results of FHT11-13. a.SPR binding curves between TNF*α* and FHT11, FHT12 and FHT13, b. Inhibition of TNF*α* activity of FHT11, FHT12 and FHT13 in cellular luciferase assay. Experiments were performed in triplicate. Error bars represent the standard deviation, c.d. Results of the qPCR cell experiments for FHT12 and FHT13, with the four tested genes being A20, CXCL2, IL6, and NK4. Data were presented as mean and SD, n.s., not significantly different, * stands for the confidence level, *: *p <* 0.05, **: *p <* 0.01, ***: *p <* 0.001, ****: *p <* 0.0001 (calculated by two way ANOVA).

## 4 Conclusion and Discussion

We have developed CYC_BUILDER, a cyclic peptide binder generation method that combines fragment assembly with reinforcement learning-assisted fragment sampling. CYC_BUILDER efficiently generates potential target binding head-to-tail or disulfide-cyclized peptides, which has been successfully applied in discovering novel TNF*α* cyclic peptide inhibitors.

Using simulation-based evaluations, CYC_BUILDER is highly effective for cyclic peptide design as currently available protein-cyclic peptide compex data is rather limited. Its performance is driven by three key features: a comprehensive tripeptide fragment library tailored for protein-protein interfaces, enabling high-affinity and diverse peptide generation; a Monte Carlo Tree Search (MCTS) framework that efficiently guides fragment sampling and peptide assembly; and advanced scoring methods to assess conformational stability, binding affinity, and cyclization propensity. Real-time decision-making is carried out without pre-training and interpretable results are provided through the tree-based structure. We conducted a 600-step performance test on the CYC-19C dataset for different fragment sampling strategies. As shown in Figure 7, the rollout method significantly impacts the convergence time and the best scores. The time-dependent method achieves the best performance. Classifying residues into seven categories (See Table S1) further facilitates better convergence. We also conducted a detailed diversity analysis for the CYC_BUILDER generated ensemble of cyclic peptide binders in TNF*α* case study to evaluate the model’s performance. The average Pairwise L-RMSD of cyclic peptides with the same length was 8.66 Å. And the average sequence similarity between the cyclic peptide ligands was 0.33. This demonstrates that CYC_BUILDER can generate diverse binding modes and sequences (See Figure S10 and Figure 4e). CYC_BUILDER runs well on a server equipped with only 10 Intel(R) Xeon(R) Gold 6132 CPUs (2.60GHz), each with 28 cores, demonstrating its ability to deliver high efficiency while requiring minimal computational resources.

We have demonstrated that CYC_BUILDER can efficiently generate cyclic peptide binders for protein targets with high structural and sequence diversity. This provides a solid foundation for applications in biomedicine, especially in protein therapeutics and targeted drug discovery. While the current version is limited to the 20 canonical amino acids, its extensible design paves the way for future enhancement, including the integration of non-canonical residues and multi-cyclic structures, enabling application potential for more diverse tasks.

## 5 Methods and Materials

### 5.1 Fragment Databases

#### Peptide fragment databases curation

The distribution of amino acid residues on the interface differs significantly from the amino acid abundance in natural proteins (Figure S3). In order to ensure that the fragments closely mimic the conformations of natural interfacial interactions, we utilized the protein-protein complex database to construct libraries of all interfacial oligopeptide fragments. X-ray crystal structures with a resolution better than 3Å from PDB (Protein Data Bank, earlier than September 10, 2021)^51^ that contain two or more chains were collected. We extracted the surface residues on the protein-protein interaction interface, which were defined as residues within 10 Å of the partner chain. After removing redundancy (aligned backbone RMSD ≥ 0.5 Å). We finally get 26,118 interface motifs. For a protein-protein complex structure, we first identified all the interfacial residues, whose atoms are within 5 Å of the partner protein, and then collected all the continuous 3 or 4 interfacial residues on sequence as the candidate tripeptide or tetrapeptide fragments. The further criteria for tripeptide selection are that satisfy *dSASA_total_* ≥ 45Å^2^ or *dSASA_central_/SASA_central_* ≥ 80% and *dSASA_central_* ≥ 30Å^2^. The criteria for 4mers are *dSASA_total_* ≥ 72Å^2^ or *dSASA_central_*_12_*/SASA_central_*_12_ ≥ 80% and *dSASA_central_*_12_ ≥ 60Å^2^ (where central refers to the central one residue of the tripeptide or the central two residues of the tetrapeptide). The redundant fragments with the same sequence and backbone RMSD ≤ 0.3Å. Finally, we obtained 1.13 million tripeptide and 0.65 million tetrapeptide fragments. The tripeptide fragment database are used for both peptide growth and the fragment-bridging cyclization. The tetrapeptide fragment database is only used for the cyclization. The distance C*β*1–C*β*2, the angles of C*α*1-C*β*1-C*β*2 and C*α*2-C*β*2-C*β*1, the dihedral of C*α*1-C*β*1-C*β*2-C*α*2 are calculated for each structure and save in the database for the geometry matching operation in the peptide cyclization step (See Figure S4 and Figure S5a for details).

#### Fragment database classification

In the expansion step, the fragment sampling was based on the feedback sampling probability. The fragments in the same class are expected to have a similar return value after sampled for peptide growth to achieve the convergence of sampling. Thus, the tripeptide fragments are classified by both the conformation and the chemical property, which are the major determination of the spliced structure as well as the stability and binding affinity score. On the other hand, the number of classes should be limited to improve the sampling efficiency. To enhance the efficiency of our model’s sampling algorithm and optimize the scale of the sampling space, we categorized the collected oligopeptides into a sampling database according to residue types (Table S1). Firstly, we classify the 20 natural amino acid residues into 7 categories based on size and polarity. Secondly, the conformation of a tripeptide (Figure S4) is described by the angles of the three C*α* atoms and the side chain orientation of the N-terminal residue and the middle residue. The side chain of the C-terminal residue is not retained in the spliced peptide, so the side chain orientation of this residue is not considered. The side chain orientation is defined as the dihedral angle between the vector of the C*α* atom to the end atom of the side chain and the plane of the three C*α* atoms, which is illustrated as the dihedrals ENDATOM1–C*α*1–C*α*2–C*α*3 or ENDATOM2–C*α*2–C*α*1–C*α*3. Each of the three angles is uniformly divided into 72 bins from 0° to 360°. We collected 3mer geometric data and classified them into 384 categories, respectively. From the initial classes, we eliminated the categories with fewer than 100 entries, resulting in a total of 112 classes of fragments for sampling. In subsequent regeneration tests, we compared the sampling efficiency and the number of convergence steps of the programs with different fragment categorization methods. The results showed that the above categorization approach, based on amino acid properties, made the sampling converge faster and resulted in cyclic peptides with higher scores (See Figure 7).

#### Disulfide-bond linked cystine pair structure database

In addition, we collected all the structures in the PDB database (before March 2021) with a resolution less than 2.5 Å, and extracted all the disulfide bond units (two linked CYS pairs). We then removed redundancy based on RMSD *<* 0.3Å, resulting in 7980 different conformations of disulfide bond structures for the disulfide ring-closure algorithm. The distance C*β*1–C*β*2, the angles of C*α*1-C*β*1-C*β*2 and C*α*2-C*β*2-C*β*1, the dihedral of C*α*1-C*β*1-C*β*2-C*α*2 are calculated for each structure and save in the database for the geometry matching operation in the peptide cyclization step. The distribution of these 4 geometry parameters are shown in Figure S5b.

### 5.2 Seed seeking algorithm

The starting seed fragment for the growth algorithm, which act as a preliminarily anchor attached to the target protein, may be directly provided by the user or generated by the seed seeking algorithm. If the complex structure of a reference peptide or protein binding with the target protein is provided, the algorithm extract the tripeptide fragments containing as many as hot-spot residues from the reference as the seeds. The hot spot residues are identified by the computational alanine-scan method in Rosetta. If only the target protein structure and the target motif on this protein were provided, the seed seeking algorithm performs a tertiary fragment search on the protein-protein interaction structure database using the MASTER method. ^52^ This approach give the backbone structure of the initial seed using the similar fragment interaction pairs in PDB database. Protein-Protein interface database described in Section 5.1 were used as the PDS(Protein Data Structure) database for MASTER method. ^52^ CYC_BUILDER treat the input target protein as the matching object and perform structural matching search on the PDS database(the cutoff is defined as an RMSD of less than 0.3 Å between the target protein and the PDS backbone structures.), and obtain a series of skeleton matched PDS motifs. Then it retrieves the corresponding PDB structure of the matched motifs, and the interacting tripeptide fragments of the partner protein, whose C*α* are all within 6 Å of any C*α* of the target motif and the center is within 7 Å of the center of the target motif are extracted as the candidate seeds (Figure S1a).^53^ The candidate seeds are scored using Equation 1, which combines the inter-molecule scoring function (*S_af_* represents the Rosetta ddG and *S_sa_* represents the interface area of the tripeptide and target protein), seed fragment orientation score(*S_ori_*) and motif matching rate (*R_mat_*). *S_ori_* evaluate the deviation of the fragment to the target motif. As is shown in Figure S1b and Equation 2, when the distance between the geometric centers of the seed fragment and the target motif is smaller than 4 Å, a penalty score is assigned based on both the distance and the orientation of the tripeptide, which is defined as the angle between the line through the two centers and the bisector of the angle formed by the 3 C*α* atoms. The matching rate score R*_mat_* refers to the ratio seed fragment residues within 5 Å of any target motif residue. The specific coefficients w_1_-w_4_ and their detailed values is in Section S1.

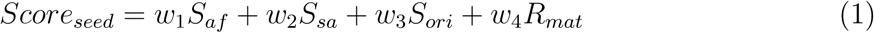

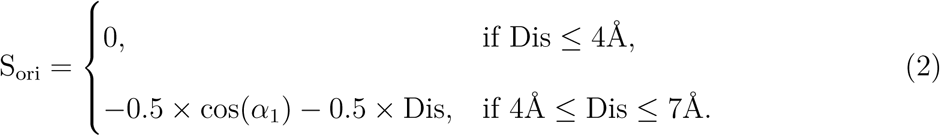

### 5.3 Splice and Cyclization methods

#### Splice algorithm for growth

The backbone of the C terminal residue of a sampled tripeptide fragment is superposed to the N terminal residue of current intermediate peptide binder (called decoy peptide), which result in the integration of the two more residues of the fragment into the current peptide. The backbone coordination of the joint residue is optimized as Equation 3, where *X*_0_ and *X*_1_ are the sets of backbone atoms (*N*, *Cα* and *C*) and *Cβ* (except for GLY) of decoy peptide and selected fragments after the superimposition, and *α* is the overlap factor. The best *α* factor is determined by resulting in a best backbone dihedral angle scoring.

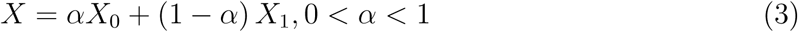

Peptides attempt to cyclization through directly head-to-tail amide bonding, disulfide linkers, and peptide fragment linkers after each fragment splicing (MCTS tree expansion).

#### Direct head-to-tail cyclization

If the distance between C*α* atoms of two terminal residues falls within the range of 3.72 to 3.92 Å, and the distance between the terminal C and N atoms is between 1.25 and 1.45 Å, an amide bond formed between the terminal atoms, which we called direct head-to-tail bonding method for cyclization.

#### Fragment based cyclization

The second ring-closure method is selecting a tripeptide or tetrapeptide fragment structure from the database (see the construction method in Section 5.5) and fill the gap between the two terminals of the decoy peptide (illustrated in Figure8b). We let Nh, C*α*h, Ch, Nt, C*α*t and Ct represent the N-terminal N C*α* C atoms and C-terminal N C*α* C atom, respectively, We calculated the distance C*α*t–C*α*h and the angles Nt-C*α*t-C*α*h and Ch-C*α*h-C*α*t, as well as the dihedral angle Nh-C*α*h-C*α*t-Ct of the decoy peptide first, and then retrieve the corresponding geometry value of the fragment in the database. The Nt, C*α*t, C*α*h, Ch atoms of the searched fragment are superimposed onto the Nt, C*α*t, C*α*h, Ch atoms of the decoy peptide, respectively, so as to bridge the two terminals of the decoy peptide. we select the side chains of the decoy peptide as the retained side chain at the two joints.

#### Disulfide bond cyclization

In the geometry matching approach of the second and third ring-closure method, the maximum disance deviation from the target value is set to 0.1 Å and the max deviation of angle or dihedral angle from the target value are set to 5° The best matching fragments are selected as the building blocks for cyclization. Subsequently, corresponding terminal residues simultaneously undergo our fragment fusion algorithm, to perform the cyclization (Figure 8b).

**Figure 7:**
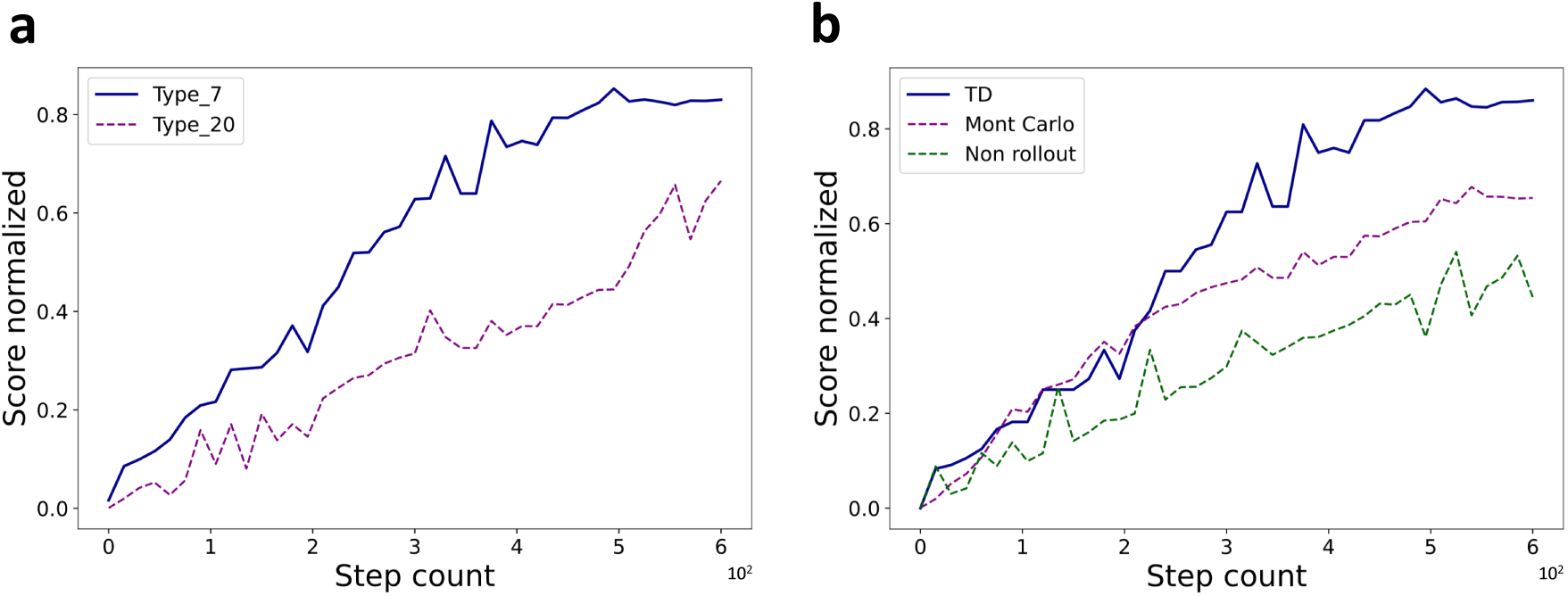
Performance analysis of CYC_BUILDER on fragment classification and rollout methods. a. Performance test on the classification criteria, Type 7 refers to the classification method used in our work and Type 20 means without residue classification, b. Performance studies of the rollout methods, TD means time-dependent method used in CYC_BUILDER, Monte Carlo is the original rollout method in MCTS and Non rollout means without rollout.

**Figure 8:**
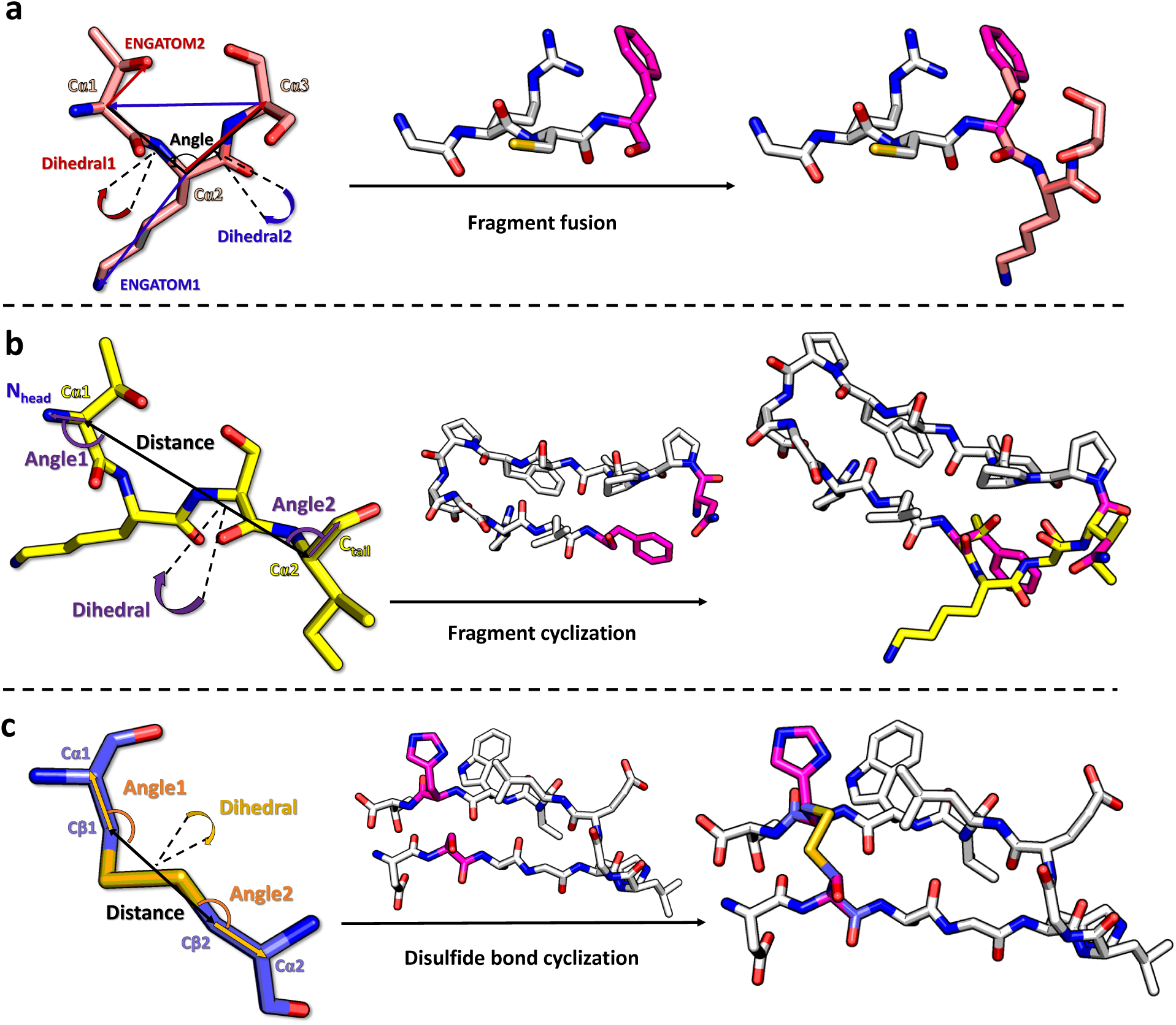
a. Schematic diagram of the fragment fusion process, b. Diagram of the fragment cyclization algorithm, c. Diagram of the disulfide bond cyclization algorithm.

#### Structure perturbation

The cyclic peptide growth process of CYC_BUILDER is semiflexible, primarily demonstrated through the model’s perturbation algorithm (see Figure S1e). CYC_BUILDER perturbs the backbone conformation of the fused residue through changing the dihedral angles *ϕandψ* by rotating 0°, ±5°, ±10°, in a manner similar to RamaProb of Rosetta.^54^ The side-chain of the residues on the newly formed fragment are repacked, using the Pymut toolkit,^55^ to eliminate intermolecular collisions and to explore more conformations. A total of at least 25 conformations are obtained and scored, and the best structure is saved as the next leaf node.

### 5.5 Soring functions

We have developed a comprehensive scoring approach that can evaluate the stability and the affinity of the decoy peptide, and also the tendency of decoy cyclization, which together with the MCTS algorithm, provide efficient guidance for growing process. Our scoring function terms are mainly divided into three categories(Equation 4), with a set of empirical weights for these terms. *S_inner_*, evaluates the conformational validity (internal collisions and the backbone torsion rationality) and the structural stability (intra-molecule hydrogen bonding). *S_inter_* estimate the binding affinity of peptide. *S_cyc_* evaluates of cyclization tendency of a peptide. The detailed terms of our score functions is in Section S1.

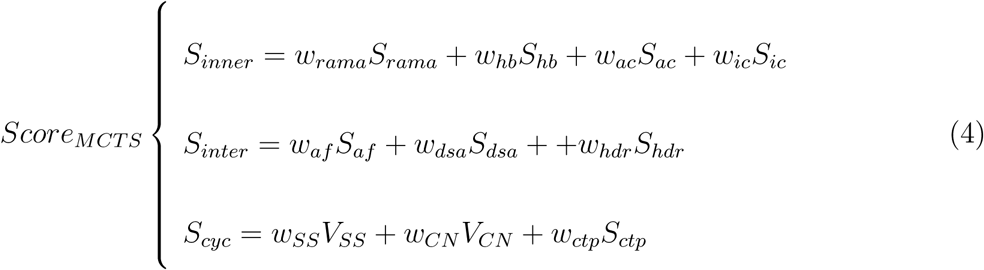

### 5.4 MCTS framework of CYC_BUILDER

The tree search algorithm in CYC_BUILDER primarily consists of the four classic steps of MCTS, namely selection, expansion, simulation, and backpropagation, and some modifications are made for the of cyclic peptide growth task. The MCTS tree is built from the seed fragment, which is the root node, and the fusion a tripeptide sampling from a fragment class means a child node linked up to the parent. In the selection step, a decoy peptide was selected by searching a best route from the root node to leaf nodes. The node selection in the route searching is based on a modification of Upper Confidence Bound for Trees (UCT)^56,57^ strategy. Each node is evaluated with Equation 6, which balance the exploitation (using the reward value) and exploration (performing a new sampling).

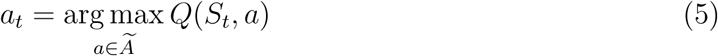

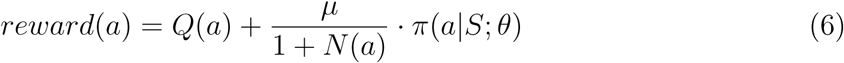

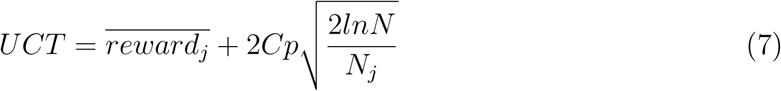

By integrating the reward and action value functions(Equation 5), the model will select the next actions for each leaf node of the next growth step. Each action selection will be evaluated by Equation 6, the reward function of CYC_BUILDER, which balance the scores and visits of the tree search process. *N* (*a*) is the number of times growth a has been visited under the current state, and *Q*(*a*) is the cumulative action value of the current state(Equation 9), *µ* is a hyper-parameter that can be adjusted, *π* is a sampling probability matrix based on the policy, *S* represents the current state (State class), and *θ* represents the inherent parameters for the probability calculation. We employ the Upper Confidence Bound for Trees (UCT) algorithm to perform state selection. This algorithm optimizes the search process by balancing exploration and exploitation, enabling a more efficient discovery of the most promising actions. As illustrated in Equation 7, 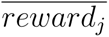 represents the average reward of node j, *N* is the visit count of the parent node, and *N_j_* is the visit count of the current node. *C_p_* is the exploration parameter, a positive real number that often needs to be adjusted based on the specific application to balance exploration and exploitation.

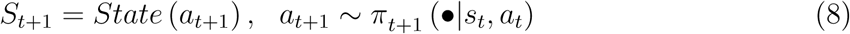

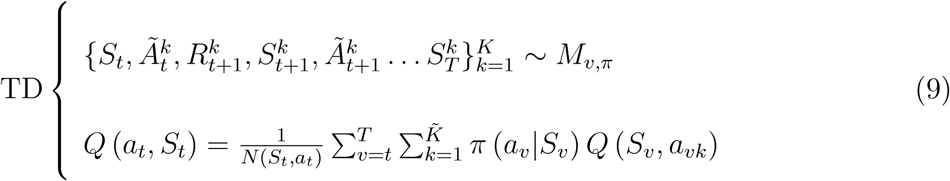

Equation 8 describes the expansion process of the model. In the expansion step, a tripeptide fragment was sampled based on the sampling probability (*π*) and splicing to the selected decoy peptide. Thus a child node was linked up to the leaf node of the selected route and this expansion was evaluated in the following simulation step.

In the simulation process, the rollout algorithm adopts a TD (time-dependent) algorithm. As shown in Equation 9, *M_v_* represents a set of Markov chains of length *T*, *K* represents the number of sampling choices. *π* denotes the sampling probability matrix, *Q* stands for the action-value function, *S* is the state and *A* is an action. This indicates that we no longer use the Monte Carlo method for roll-out. Instead, we evaluate the action value of the current node by extending one layer of leaf nodes downward (*T* = *t* + 1) and compute a weighted average of the action values of all possible leaf nodes using the sampling probability *π*.

Based on the results of simulation and backpropagation, according to Equation 9, the action-value *Q*(*s, a*) is updated through total reward and *N* . The sampling policy of each node is updated according to all state values, as shown in Equation 11.

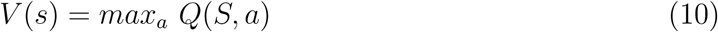

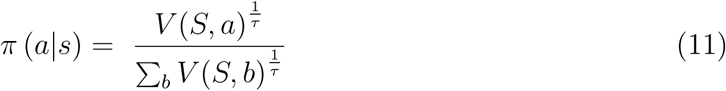

*V* represents the state value which is defined as the highest expected return achievable by taking the best possible action from all available options starting from that state(Equation 10). a and b are actions, *τ* represents the temperature parameter. Similar to AlphaZero,^58^ a larger *τ* means smaller the difference between different states, and a greater proportion of exploration. To achieve greater diversity, the model’s *τ* value is set to 1.2.

### 5.6 Experimental methods for testing the activity of TNF***α*** cyclic peptide binders

#### Expression and purification of human wild-type and mutant human TNF*α*

Firstly, high purity TNF*α* protein was prepared following the published procedures.^59^ Then, We mutated the wild-type TNF*α* plasmid using the Mut Express II Fast Mutagenesis Kit V2 (Vazyme) and the experimental protocol provided by the manufacturer to obtain the TNF*α* mutant plasmid. We purified mutant TNF*α* proteins using the same buffers and procedures as those used for wtTNF.

#### Surface plasmon resonance (SPR)

Binding kinetics analysis was performed using the Biacore T200 system (GE Healthcare) at 25^◦^C, and all experiments were performed in HBS-P buffer (10 mM HEPES, 150 mM NaCl, 0.05% surfactant, pH 7.4) with a flow rate of 30 *µ*l/min.. TNF*α* protein was immobilized on a CM5 chip by amino coupling to a response of 3600 resonance units (RUs). The reference channel was similarly activated and blocked without TNF*α* protein. The cyclic peptides were dissolved in HBS-P buffer, and different concentrations of cyclic peptides were serially injected into the experimental channel to detect binding parameters with 120 s binding and 240 s dissociation per cycle. Injection of 5 mM NaOH regenerated the chip after each sample flow-through. The obtained data were globally fitted using a steady-state model provided by Biacore BIAevaluation 4.1 software.

#### Luciferase assay

The inhibitory activity of cyclic peptides against TNF*α* was detected by cellular assay as described previously.^60^ Briefly, HEK293T cells were grown to 80% confluence in 6-well cell culture plates at 37^◦^C in DMEM medium (Gibco) containing 10% FBS (Gibco). Then, the cells were transfected with purified 2.4 *µ*g pGL4.32 (luc2P/NF-*κ*B-RE/Hygro) plasmid and 1.6 *µ*g pGL4.74 (hRluc/TK) plasmid with PEI transfection reagent (Proteintech, PR40001). After 24 h, the transfected cells were plated in a 96-well plate at a density of 40,000 cells per well. Simultaneously, TNF*α* was mixed with different concentrations of the cyclic peptides and incubated at 37^◦^C for 12 h. Then, the mixtures were added to each well respectively and stimulated the cells for 6 h. Equal amounts of TNF*α* without the cyclic peptides were added to the cells as negative control. Fluorescence intensity was measured using the Dual-Glo Luciferase Assay System (Promega) with a BioTek Synergy 4 Multi-Mode Microplate Reader. All results were calculated from three replicates. The final concentration of TNF*α* in each well is 1 ng/mL (prepared in DMEM with 1 mg/mL BSA).

#### Quantitative real-time PCR (RT-qPCR)

HEK293T cells were stimulated using 4 ng/mL of wild-type TNF*α* or a mixture of 4 ng/mL TNF*α* with different cyclic peptides pre-incubated at 37°C for 12 h was added to each well of a 6-well cell culture plates. Cells were collected and lysed at the corresponding time points (1, 2, 3, 6, 10 h). Total RNA of each sample was extracted from previously processed cells using the FastPure Cell/Tissue Total RNA Isolation Kit V2 (Vazyme) according to the manufacturer’s instructions. Then, HiScript III RT SuperMix for qPCR (+gDNA wiper) kit (Vazyme) was used to reverse transcribe total RNA to the corresponding cDNA.

To determine the expression level of the four selected genes, including A20, CXCL2, IL6 and NK4, four pairs of primers were designed. The qPCR reaction was performed using the following forward (F) and reverse (R) primers:

For A20: F: 5^′^−CACGCTCAAGGAAACAGACA−3^′^, R: 5^′^−CACGCTCAAGGAAACAGACA−3^′^; For CXCL2: F: 5^′^−TGCCAGTGCTTGCAGAC−3^′^, R: 5^′^−TCTTAACCATGGGCGATGC−3^′^; For IL6: F: 5^′^−ATGCCCATCACTCGGATGC−3^′^, R: 5^′^−CCCTGCTTTGTATCGGCCTG−3^′^; For NK4: F: 5^′^−AAAATGCAAAATGCAGAATCAGG−3^′^, R: 5^′^−TAAGCCGCCACTGTCTCCAG−3^′^.

The data were analyzed using the 2^−ΔΔ^*^CT^* method to calculate the fold change of the target genes. The expression levels of the treated target genes were normalized to the expression level of the endogenous reference gene GAPDH at 0 hours.

#### CD spectrum

Wild-type and mutant TNF*α* proteins were solubilized in 20 mM K*H*_2_P*O*_4_/*K*_2_HP*O*_4_ (pH 7.4) to a final concentration of 0.2 mg/ml. CD spectra in the far UV region (190-260 nm) were measured using a MOS 450 AF/CD (Biologic, France) with a 1 mm quartz cuvette at 25°C. The average of three cumulative measurements was taken and smoothed using the standard noise reduction technique provided with the instrument.

## Acknowledgement

This work was supported by the National Natural Science Foundation of China (21977007, 32101003) and the National Key R&D Program of China (2022YFA1303700).

## Supporting Information Available

### Supporting information

The rest of the introduction to the model scoring function, database details, further introduction to redock results, ablation experiments, MD parameter settings and details of MMPBSA results and additional wet experiment results.

### Data and code availability

Code and data of our work will be available upon publication.

