## Supplemental information for "Target-based *de novo* design of cyclic peptide binders"

### 1 Algortitms and score functions

#### Details of seed seeking algorithms

The specific parameter settings of seed seeking algorithms are as follows:

$$\begin{cases} w_1 = 0.4, \mathrm{Dis} \leq 4\text{Å} \\ w_2 = 0.2, \mathrm{Dis} \leq 4\text{Å} \\ w_3 = 0, \mathrm{Dis} \leq 4\text{Å}; 0.2, Dis \geq 4\text{Å} \\ w_4 = 0.2, \mathrm{Dis} \leq 4\text{Å} \end{cases} \tag{1}$$

As illustrated in Figure S1 and Equation 1, when the seed fragment falls within the Core region, we do not calculate its affinity score, setting $w_3$ to 0. In the Peripheral region, we enable all scoring terms and see flowchart in Figure S1 for details.

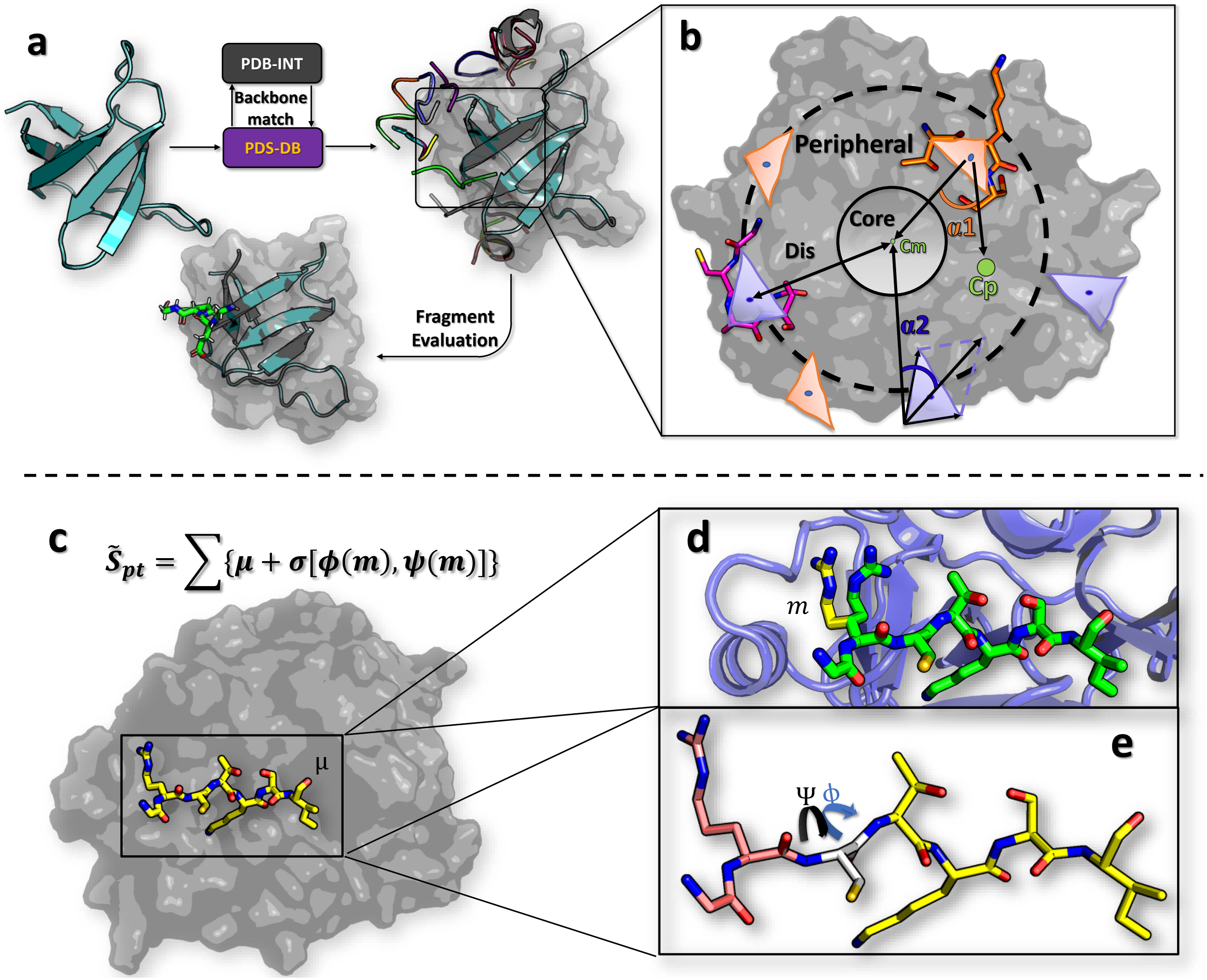


Figure S1: a. Schematic diagram of the Seed search algorithm, homologous fragments on the protein-protein interaction interfaces with similar skeletons are found from the natural PDB database and are simply optimized, serving as the root nodes for the MCTS search algorithm. b.The figure illustrates the scoring method for the orientation of the seed, **Cm** and **Cp** refers to the C$\alpha$ geometric center of the binding motif and protein, respectively. **Core** is the area within 4 Å of Cm, and **Sphere** is the area with a distance between 4-7 Å away from Cm. **Dis** refers to the distance from the geometric center of the fragment to Cm. $\alpha$**1** is the angle formed by the Cp, the geometric center of the fragment, and Cm. $\alpha$**2** is the angle between the bisector of the fragment's C$\alpha$s angle and the line connecting the C$\alpha$ at the middle position of the fragment with Cm, c. Schematic diagram of the complex structure. The complex structure we obtained is an ensemble composed of the ligand structures obtained through perturbation and the receptor, d. Schematic diagram of side chain repacking, where m represents a type of rotamer, e. Schematic diagram of main chain perturbation, where psi and phi respectively represent the two types of main chain dihedral angles undergoing perturbation.

#### Details of score functions

$S_{inner}$: We initially adopted a stepwise function $S_{hb}$ based on the statistical distribution of geometric parameters in Rosetta[1] (see Figure S1) to quickly score intramolecular hydrogen bonds. Additionally, Rosetta's statistical potential of backbone dihedral angles was utilized as $S_{rama}$.[1] Residue composition score ($S_{ac}$, Equation S2) was designed to constrained the number of prolines($Rt_{pro}$), hydrophobic residues($Rt_{np}$) and aromatic residues($Rt_{aro}$) to be around $r1 = 2$, $r2 = 0.3 *$ (number of all residues) and $r3 = 0.1 *$ (number of all residues) by default. Prolines can enhance the rigidity of the backbone[2] and an appropriate ratio of hydrophobic and hydrophilic residues is a simple way to control the solubility and logP of the designed peptides. $S_{ic}$ is a penalty score for intramolecular collisions. The number of side-chain-related collisions ($sc_{coll}$ ), and collisions between backbone atoms ($bb_{coll}$) are checked first with a definition of the distance of the two atoms smaller than 0.7 times the sum of the van der Waals radii, and then calculated the collision score with Equation S3. Collisions involving the backbone incur a higher penalty and are normalized using a sigmoid function.

$$S_{ac} = -(|Rt_{pro} - r1| + |Rt_{np} - r2| + |N_{aro} - 2|) \tag{2}$$

$$S_{ic} \ = \ -Sigmoid[(\sum sc_{coll} \ + \ 5 \sum bb_{coll})] \tag{3}$$

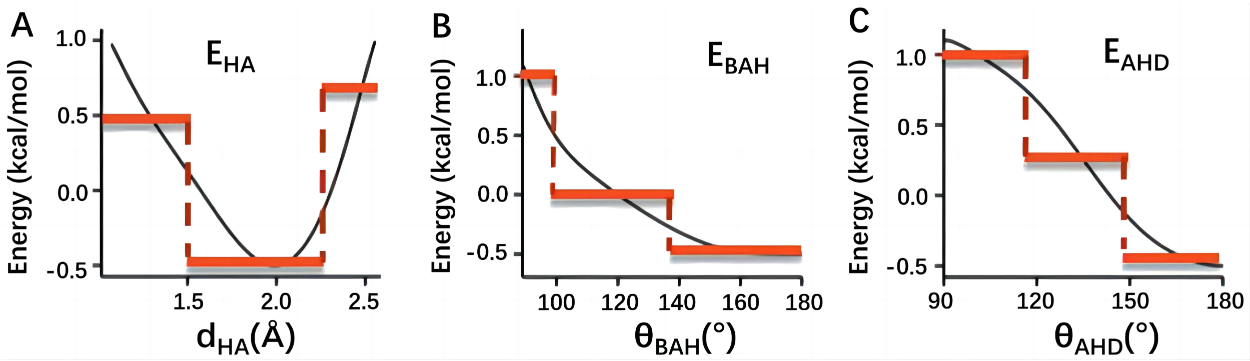


Figure S2: A,B,C represents the step-wise potential of CYC_Builder for $E_{HA}$, $E_{BAH}$ and $E_{AHD}$

Figure S2 shows the step-wise hydrogen bond potential of CYC_Builder, derived from Rosetta's statistical potential.[1] H stands for hydrogen bond donors, A represents hydrogen bond acceptors, and D denotes donors. $d_{HA}$, $\theta_{BAH}$, and $\theta_{AHD}$ represent the distance and angles among B, H, and A, respectively. The energy $E_{dHA}$ is set to 0.5, -0.5, and 0.75 for the $d_{HA}$ bins of ≤1.5Å, 1.5Å≤d≤2.25Å, and ≥2.25Å, respectively. The $E_{BAH}$ is set to 1.0, 0.0, and -0.5 for the $\theta_{BAH}$ bins of ≤100°, 100°-140°, and ≥140°, respectively. The $E_{AHD}$ is set to 1.0, 0.25, and -0.5 for the $\theta_{AHD}$ bins of ≤120°, 120°-150°, and ≥150°, respectively. The scoring function is set as shown in Equation S4

$$S_{hb} = -\sum (E_{HA} + 1.5E_{BAH} + E_{AHD}) \tag{4}$$

$S_{inter}$: Peptide–target protein interaction score estimate the binding affinity using the 3D grid-based SDOCK score for the efficient calculation,[3] Including Van der Waals energy $E_{LJ}$, electrostatic energy $E_e$, desolvation energy $E_{LK}$, and and geometric collision $E_c$. These energy terms are all expressed as the product of the atom property and the field strength. The field strength values for a point near the targeting site were pre-calculated and assigned to the grid point and the total binding energy for a decoy peptide can be quickly calculated with Equation S5. The center of the target motif is set as the center of the grid and the size is set to cubic with a 30 Å side length. In order to improve the accuracy of scoring, we used a much finer grid (grid space is 0.6 Å) compared to the unbound docking program used (grid space is 1.2 Å) . We use the per residue interaction score to evaluate the binding complementary and control the length of the product cyclic peptide $S_{af}$. The buried surface area $S_{dsa}$, is calculated as the reduction of the total Solvent Accessible Surface Area ($dSASA$) caused by the protein-peptide binding, as shown in Equation S6. We expect to control the total number of hydrophobic residues of the designed peptide so as to have a good solubility, and also expect the hydrophobic residues are most buried in the complimentary hydrophobic interface. Thus, hydrophobic exposure ratio, that is the total $SASA$ of a hydrophobic residue in the free state minus its $SASA$ buried in the complex divided by its total $SASA$, greater than 40% and an exposed $SASA$ exceeding 80 Å$^2$ will incur penalties as described in Equation S7.

$$\Delta E_{SDOCK} = w_{LJ}E_{LJ} + w_e E_e + w_{LK}E_{LK} + w_c E_c \tag{5}$$

$$S_{dsa} = Sigmoid(\frac{dSASA}{800}) \tag{6}$$

$$S_{hdr} = -\frac{N_{exposed}}{N_{peptide}} \tag{7}$$

**$S_{cyc}$:** The cyclic potential developed in ADCP program was employed to evaluates of cyclization tendency of a decoy peptide (Equation S8). When the distance between the C$\alpha$ atoms at both ends (dC$\alpha$) exceeds 5 Å, scoring is based on dC$\alpha$ to encourage a decreasing distance between the ends. Once the ends are sufficiently close, the distance between the terminal C and N atoms (dCN) is incorporated as a criterion, with quadratic constraints applied to dC$\alpha$ and dCN at standard values of 3.819 Å and 1.345 Å, respectively. In Equation S8, $T_{ctd1} = 3.819$ and $T_{ctd2} = 3.819^2 + 1.345^2$. The potential $V_{CN}$ is normalized by a sigmoid function. $V_{SS}$ is the is the number of the matched cysteine pair structure from the disulfide database, which represent the statistical potential of the disulfide bond conformation. $S_{ctp}$

is the cumulative sum of $C\alpha$ distances of peptide subtract the distance between the first and last $C\alpha$s(Equation S9), which is another method to estimate the cyclization tendency. A smaller Sctp means a low tendency for cyclization.

$$V_{CN} = \begin{cases} Sigmoid\left(-\frac{dC\alpha}{T_{ctd1}}\right), \ dC\alpha > 5\ \text{Å} \\ Sigmoid[-\frac{(dC\alpha\ -\ 3.819)^2\ +\ (dCN\ -\ 1.345)^2}{T_{ctd2}}], dC\alpha \leq 5\ \text{Å} \end{cases} \tag{8}$$

$$S_{ctp} = Sigmoid\left[\frac{\Sigma_{i=1}^{N_{peptide}} d_i - D}{N_{peptide}}\right] \tag{9}$$

Based on Equation 11, all the coefficients for the scoring function in the MCTS process are as follows:

$$\mathbf{W}_{MCTS} \begin{cases} w_{rama} = 0.05, w_{hb} = 0.1, w_{ac} = 0.05, w_{ic} = 0.05 \\ w_{af} = 0.2, w_{dsa} = 0.2, w_{hdr} = 0.05 \\ w_{SS} = 0.1, w_{CN} = 0.1, w_{ctp} = 0.1 \end{cases} \tag{10}$$

### 2 Fragment geometric parameters

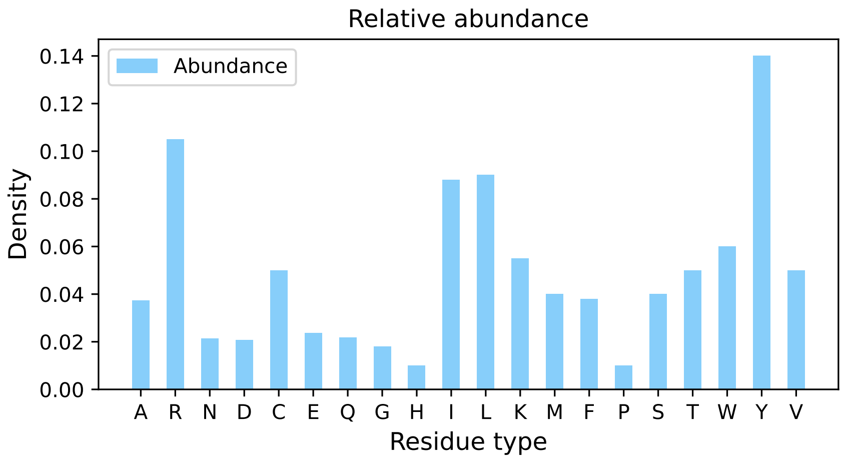


Figure S3: Distribution of amino acid residues on the protein-protein interaction interface, normalized according to the abundance of natural protein amino acids.

Table S1: Sampling residue type classification

| **Type** | **Residue Name** | **Side chain** |
|---|---|---|
| Small non-polar | Gly/G | $-H$ |
| | Ala/A | $-CH_3$ |
| | Pro/P | $-C_3H_6$ |
| | Val/V | $-CH-(CH_3)_2$ |
| Small polar | Ser/S | $-CH_2OH$ |
| | Cys/C | $-CH_2SH$ |
| | Thr/T | $-CH_2(CH_3)-OH$ |
| Medium non-polar | Ile/I | $-CH(CH_3)-CH_2-CH_3$ |
| | Leu/L | $-CH_2-CH(CH_3)_2$ |
| | Met/M | $-(CH_2)2-S-CH_3$ |
| Negatively charged | Glu/E | $-(CH_2)_2-COOH$ |
| | Asp/D | $-CH_2-COOH$ |
| Medium polar | Gln/Q | $-(CH_2)_2-CONH_2$ |
| | Asn/N | $-CH_2-CONH_2$ |
| | His/H | $-CH_2-C_3H_3N_2$ |
| Positively charged | Lys/K | $-(CH_2)_4-NH_2$ |
| | Arg/R | $-(CH_2)_3-NHC(NH)NH_2$ |
| Aromatic | Phe/F | $-CH_2-C_6H_5$ |
| | Tyr/Y | $-CH_2-C_6H_4-OH$ |
| | Trp/W | $-CH_2-C_8NH_6$ |

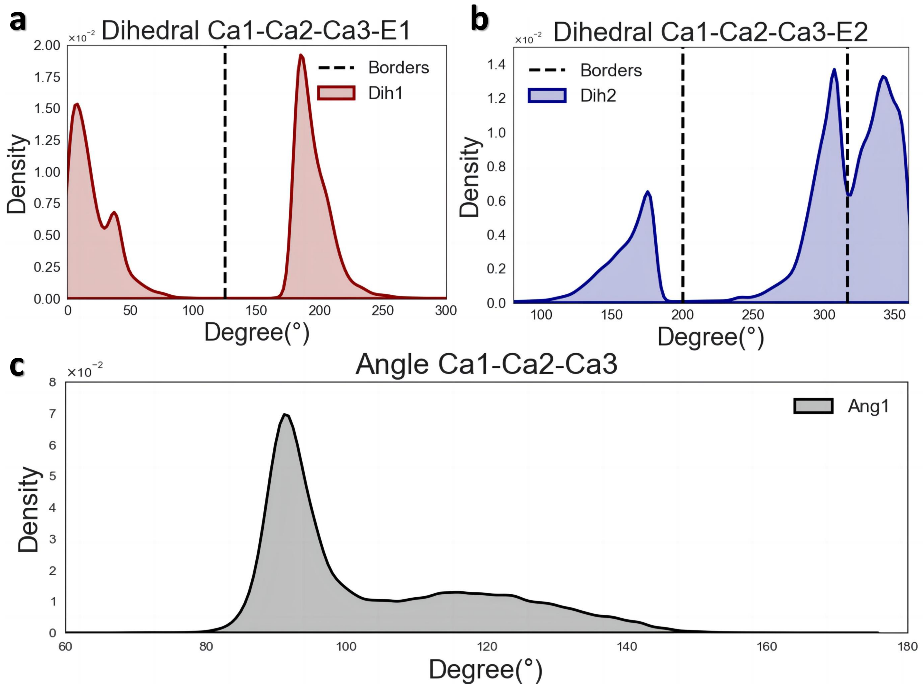


Figure S4: a. Diheral distribution of $C\alpha1 - C\alpha2 - C\alpha3 - ENDATOM1$, b. Diheral distribution of $C\alpha1 - C\alpha2 - C\alpha3 - ENDATOM2$, c. Angle distribution of $C\alpha1 - C\alpha2 - C\alpha3$

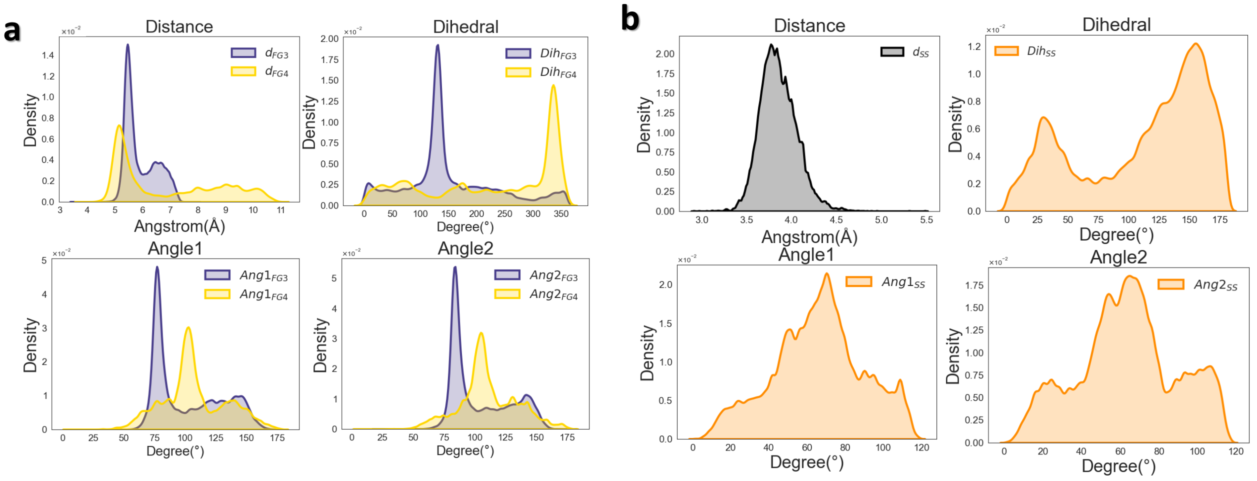


Figure S5: a. Geometric parameters distribution of 3mer and 4mer cyclization fragments, b. Geometric parameter distributions of disulfide bond building blocks.

We determined the range of geometric parameters for the model-matching fragments based on these binding parameters, and grouped the different geometric parameters using bins spaced at 10° and 0.1 Å.

$$FG3\begin{cases} d_{FG3}\left(\mathrm{C}\alpha_{head}-\mathrm{C}\alpha_{tail}\right)\in(3.5\text{ Å},8.0\text{ Å})\\ Angle1_{FG3}\left(\mathrm{C}\alpha_{head}-N_{head}-C_{tail}\right)\in(0^{\circ},180^{\circ})\\ Angle2_{FG3}\left(N_{head}-C_{tail}-\mathrm{C}\alpha_{tail}\right)\in(20^{\circ},180^{\circ})\\ \mathrm{Dihedral}_{FG3}\left(\mathrm{C}\alpha_{head}-N_{head}-C_{tail}-\mathrm{C}\alpha_{tail}\right)\in(10^{\circ},360^{\circ})\end{cases}\tag{11}$$

$$FG4\begin{cases} d_{FG4}\left(\mathrm{C}\alpha_{head}-\mathrm{C}\alpha_{tail}\right)\in(3.8\text{ Å},15.6\text{ Å})\\ Angle1_{FG4}\left(\mathrm{C}\alpha_{head}-N_{head}-C_{tail}\right)\in(0^{\circ},180^{\circ})\\ Angle2_{FG4}\left(N_{head}-C_{tail}-\mathrm{C}\alpha_{tail}\right)\in(0^{\circ},180^{\circ})\\ \mathrm{Dihedral}_{FG4}\left(\mathrm{C}\alpha_{head}-N_{head}-C_{tail}-\mathrm{C}\alpha_{tail}\right)\in(0^{\circ},360^{\circ})\end{cases}\tag{12}$$

$$SS\begin{cases} d_{SS}\left(\mathrm{C}\beta 1-\mathrm{C}\beta 2\right)\in(3.3\text{ Å},4.5\text{ Å})\\ Angle1_{SS}\left(\mathrm{C}\alpha 1-C\beta 1-C\beta 2\right)\in(78^{\circ},95^{\circ})\\ Angle2_{SS}\left(C\beta 1-C\beta 2-\mathrm{C}\alpha 2\right)\in(78^{\circ},95^{\circ})\\ \mathrm{Dihedral}_{SS}\left(\mathrm{C}\alpha 1-C\beta 1-C\beta 2-\mathrm{C}\alpha 2\right)\in(20^{\circ},90^{\circ})\end{cases}\tag{13}$$

### 3 Additional Redock Analysis

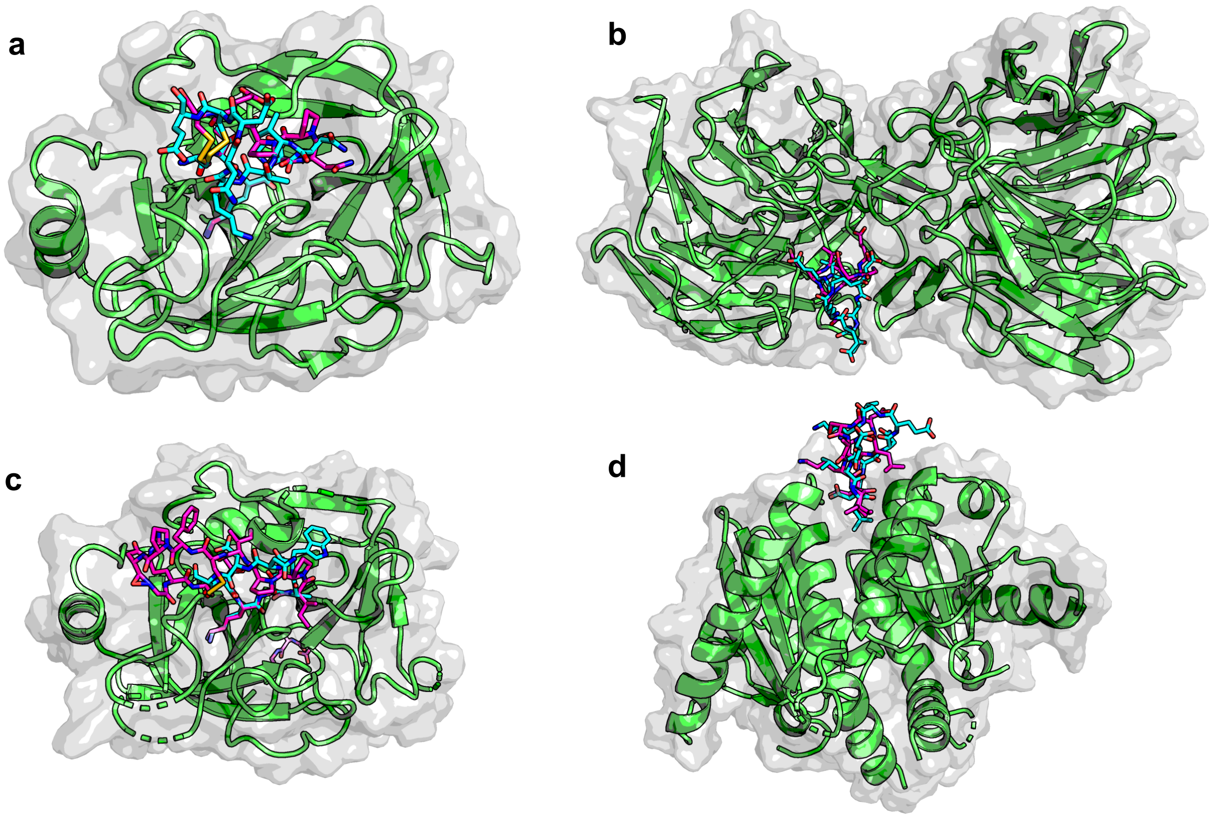


Figure S6: Representative structures from regeneration tests, with indigo for generated ligands and magenta for original ligands: (a) PDB ID 1SMF complex, linked by a disulfide bond ring; (b) PDB ID 3ZGC complex, linked by an amide bond ring; (c) PDB ID 3P8F complex, linked by an amide bond ring; (d) PDB ID 3AV9 complex, linked by an amide bond ring.

Table S2: Comparison of Amide and Disulfide Data

| **PDB (Amide)** | **BB-BB**$_{t100p}$ | **BB-SC**$_{t100p}$ | **PDB (Disulfide)** | **BB-BB**$_{t100p}$ | **BB-SC**$_{t100p}$ |
|---|---|---|---|---|---|
| 1sfi | 0.33 | 0.18 | 1smf | 0.23 | 0.43 |
| 3av9 | 0.5 | 0.31 | 1vwd | 0.37 | 0.21 |
| 3p8f | 0.44 | 0.22 | 2ck0 | 0.18 | 0.09 |
| 3zgc | 0.42 | 0.09 | 3g5v | 0.24 | 0.24 |
| 5xn3 | 0.31 | 0.12 | 3p72 | 0.22 | 0.08 |

| **PDB (Amide)** | **BB-BB$_{t100p}$** | **BB-SC$_{t100p}$** | **PDB (Disulfide)** | **BB-BB$_{t100p}$** | **BB-SC$_{t100p}$** |
|---|---|---|---|---|---|
| 4k1e | 0.19 | 0.09 | 3wnf | 0.42 | 0.23 |
| 4kel | 0.2 | 0.15 | 4ib5 | 0.31 | 0.21 |
| | | | 4m1d | 0.37 | 0.25 |
| | | | 5djc | 0.34 | 0.3 |
| | | | 5eoc | 0.34 | 0.19 |
| | | | 5th2 | 0.16 | 0.13 |
| | | | 5vb9 | 0.28 | 0.37 |

### 4 MD & MMPBSA settings and additional *de novo* generation information

Molecular Dynamics simulations were performed using GROMACS 2023[4] with the CHARMM36 forcefield.[5] Each peptide or complex was dissolved in cubic box of explicit TIP3P waters[6] and neutralized with either sodium or chloride ions. The solvated systems were energy-minimized using the steepest descent minimization method. Next, the system was equilibrated for 10 ns under the NVT ensemble with position restraints (1000 kJ/mol$^{-1}$nm$^{-1}$) applied on all the heavy atoms of the peptide. During this equilibration, pressure coupling to 1 atm was performed with the Berendsen barostat,[7] and temperature coupling to 310.5 K using the velocity-rescaling thermostat.[8] From each equilibrated system, simulations of 100 ns were performed in the NPT ensemble. The systems were simulated using periodic boundary conditions. A cutoff at 10 Å was used for van der Waals and short-range electrostatic interactions. The Particle-Mesh Ewald (PME) summation method was used for the long-range electrostatic interactions.[9] The Verlet cutoff scheme was used.[10] All chemical bonds were constrained using the LINCS algorithm.[11] The integration time-step was 2 fs, and simula-

Table S3: Detailed information of CYCB-19C dataset.

| PDB_ID | Receptor name | Bio-activity | Reference DOI |
|---|---|---|---|
| **1sfi (CN)** | Bovine $\beta$-trypsin | Ki=0.1nM | 10.1006/jmbi.1999.2891 |
| **3av9 (CN)** | Integrase(C56S, F139D, F185H mutation) | IC50=435uM | 10.1002/cbic.201100350 |
| **3zgc (CN)** | Kelch-like Ec-associated Protein 1 | - | 10.1107/S174430911301124X |
| **4k1e (CN)** | Kallikrein-related peptidase 4 (KLK4) | Ki=3.59nM | 10.1038/srep35385 |
| **4kel (CN)** | Human Kallikrein-4 | Ki=0.0387nM | 10.1021/acs.biochem.9b00191 |
| **5xn3 (CN)** | SPRY domain-containing SOCS box protein 2 | Kd=671nM | 10.1016/j.bbrc.2017.05.122 |
| **1smf (SS)** | Trypsin | Ki=0.12uM | 10.1093/oxfordjournals.jbchem.a124491 |
| **1vwd (SS)** | Sreptavidin | 230nM | 10.1074/jbc.272.20.13220 |
| **2ck0 (SS)** | Immunoglobulin, light chain | - | To be published |
| **3g5v (SS)** | mAb806 Antibody targeting a locally misfolded region of tumor associated EGFR | Kd=16nM | 10.1073/pnas.0811559106 |
| **3p72 (SS)** | Platelet glycoprotein Ib alpha chain | - | 10.1182/blood-2009-05-224170 |
| **3wnf (SS)** | Gag-Pol polyprotein | - | To be published |
| **4ib5 (SS)** | Casein kinase II subunit alpha | Kd=559.67nM | 10.1021/cb3007133 |
| **4m1d (SS)** | Fab mAb 447-52D Light Chain | Kd=23nM | 10.1021/bi400645e |
| **5djc (SS)** | Ig gamma-1 chain C region | - | 10.1016/j.str.2016.02.013 |
| **5eoc (SS)** | Fab C2 | Kd=30.3nM | https://doi.org/10.1128/JVI.02397-15 |
| **5th2 (SS)** | Cetuximab Fab | Kd=30uM | 10.1107/S2053230X16015612 |
| **5vb9 (SS)** | Interleukin-17A | Kd=1.3uM | 10.1371/journal.pone.0190850 |

tions were analyzed using GROMACS tools. We calculated the root-mean-square deviation (RMSD) of the position of the C$\alpha$ atoms of the peptides, compared to the initial conformation, using *gmx rms*.

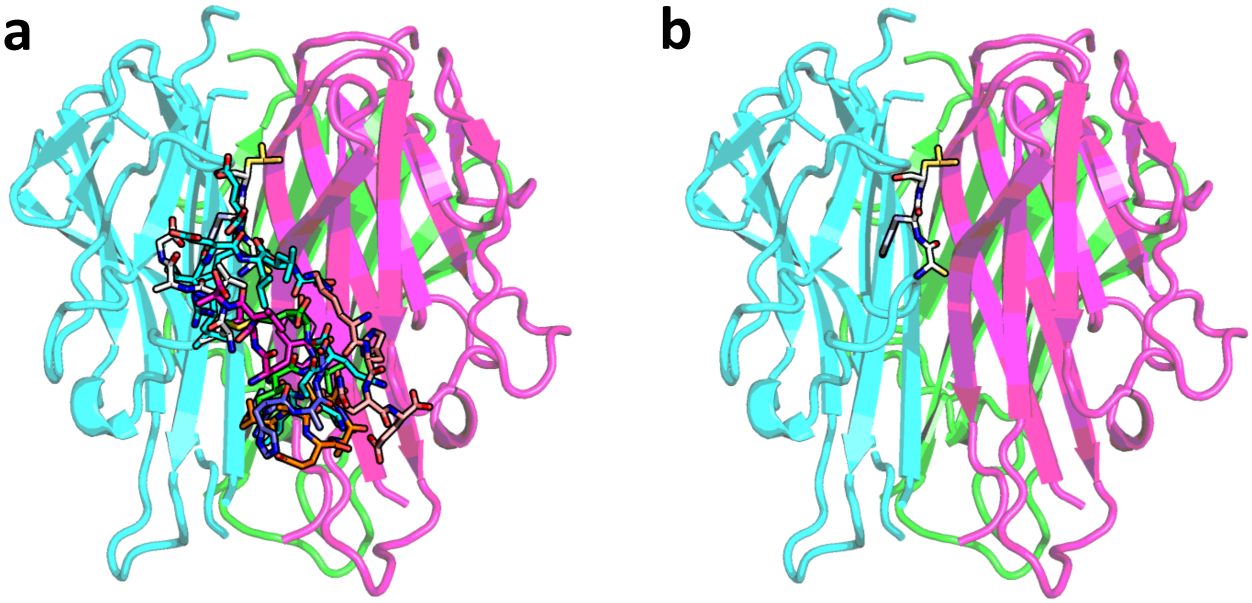


Figure S7: a.Complex structure of TNF$\alpha$ and seeds for the *de novo* generation test, b.complex structure of Seed$_{fht03}$

The MMPBSA analysis were established by gmx_MMPBSA (version 1.6.1).[12] The molecular mechanics/Poisson-Boltzmann surface area method (recommended for the CHARMM force field) was used. Single-term total non-polar solvation free energy($inp = 1$) was used. The molecular surface was used for cavity term calculation ($use\_uav = 0$). The charmm_radii ($PBRadii = 7$) was used to build amber topology files. Default parameters were applied for other terms.

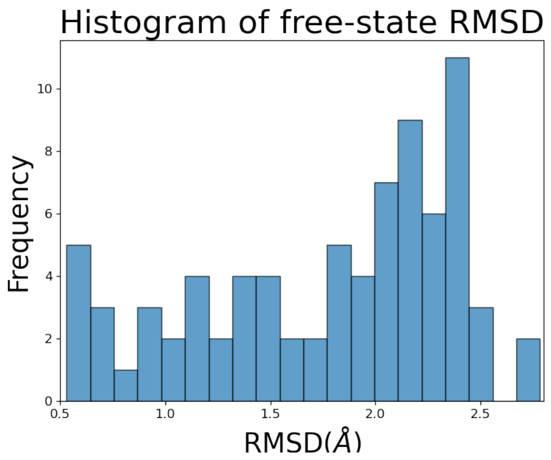


Figure S8: Bar chart of the RMSD distribution of the trajectories for the free-state cyclic peptide simulation.

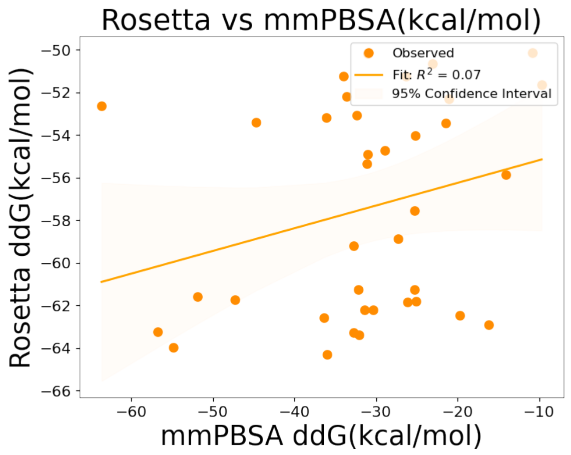


Figure S9: Linear correlation analysis between the ddG values calculated by Rosetta and those obtained from MMPBSA.

Table S4: MMPBSA Decomposition Results of FHT03

| **Chain** | **Name** | **MMPBSA ddG(kcal/mol)** |
|---|---|---|
| | VAL17 | -0.82 |
| | ALA18 | 0.38 |
| | ASN19 | -0.05 |
| | PRO20 | -0.69 |

| **Chain** | **Name** | **MMPBSA ddG(kcal/mol)** |
|---|---|---|
| | GLN21 | -0.10 |
| | LEU29 | -0.32 |
| | ASN30 | -0.06 |
| | **ARG31** | **-20.92** |
| | **ARG32** | **-9.94** |
| | ALA33 | -2.50 |
| | ASN34 | -1.44 |
| | ALA35 | -0.50 |
| | SER147 | 0.25 |
| | GLY148 | -0.71 |
| | ARG82 | 1.38 |
| | ALA84 | -0.15 |
| | SER86 | 0.05 |
| | **TYR87** | **-4.68** |
| | GLN88 | -0.03 |
| | THR89 | -0.27 |
| Receptor:C | LYS90 | -0.24 |
| | VAL91 | -2.88 |
| | ASN92 | -0.37 |
| | LEU93 | 0.12 |
| | GLN125 | -0.11 |
| | LEU126 | 0.01 |
| | GLU127 | 0.34 |
| | ASP130 | -0.51 |
| | ALA1 | -1.42 |

| Chain | Name | MMPBSA ddG(kcal/mol) |
|---|---|---|
| | TYR2 | -3.27 |
| | LEU3 | -2.50 |
| | VAL4 | -0.81 |
| | LEU5 | -1.63 |
| | GLU6 | -7.60 |
| | LEU7 | -0.25 |
| | ASP9 | -3.98 |
| | PRO10 | -0.55 |
| | LEU11 | -2.67 |

### 5 Diversity analysis of Case TNF-$\alpha$

We further analyzed the performance of the model in terms of diversity on the TNF-alpha case, and found that the cyclic peptides generated by the model were capable of binding to diverse sites. The average Pairwise L-RMSD of cyclic peptides with the same length was 8.66 Å. Additionally, using the pairwise2 module of Biopython,[13] the average sequence similarity between cyclic peptide ligands was calculated to be 0.33. This demonstrates that the model can generate diverse scaffolds and binding modes, and exhibits a good level of sequence diversity.

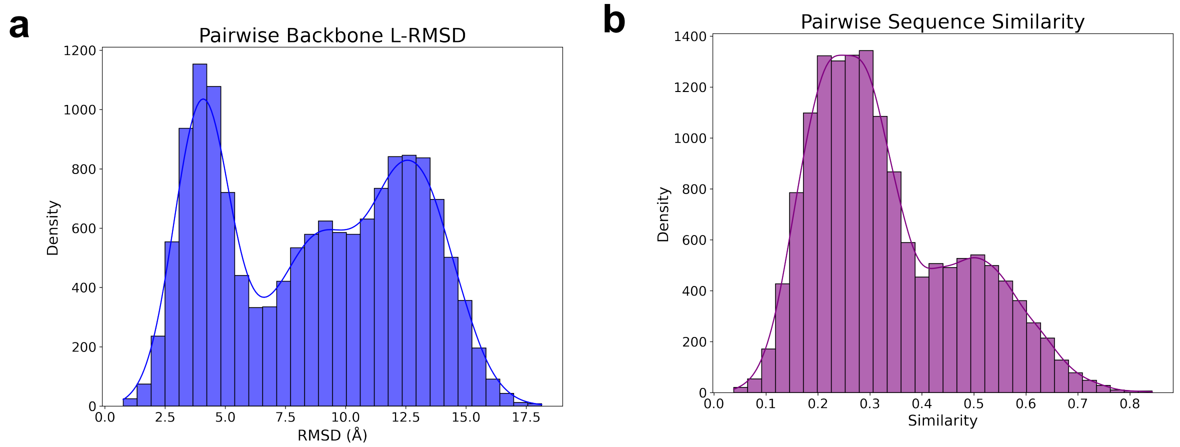


Figure S10: a.Pair-wise L-RMSD distribution of generated cyclic peptide binders, b. Pair-wise sequence similarity distribution of them.

### 6 Additional experimental data

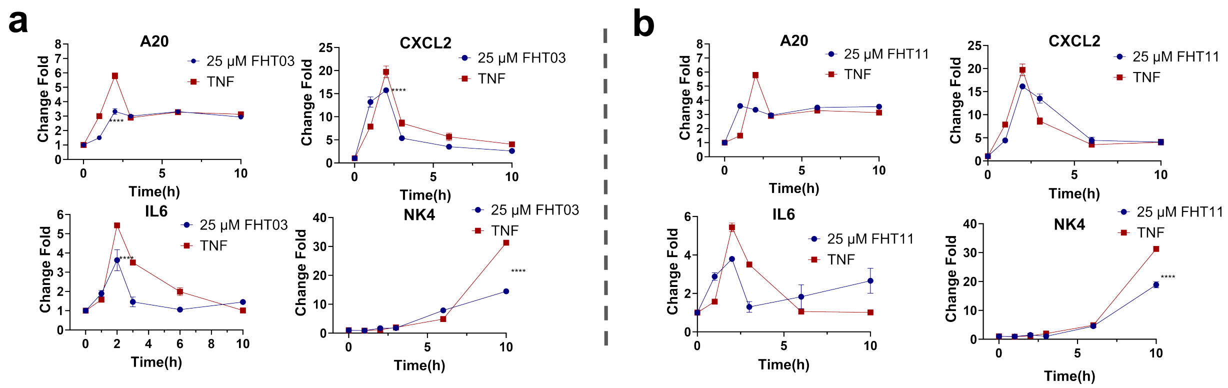


Figure S11: Results of the qPCR cell experiments for other ligands, with the four tested genes being A20, CXCL2, IL6, and NK4. a,b represents FHT03, FHT11, respectively.

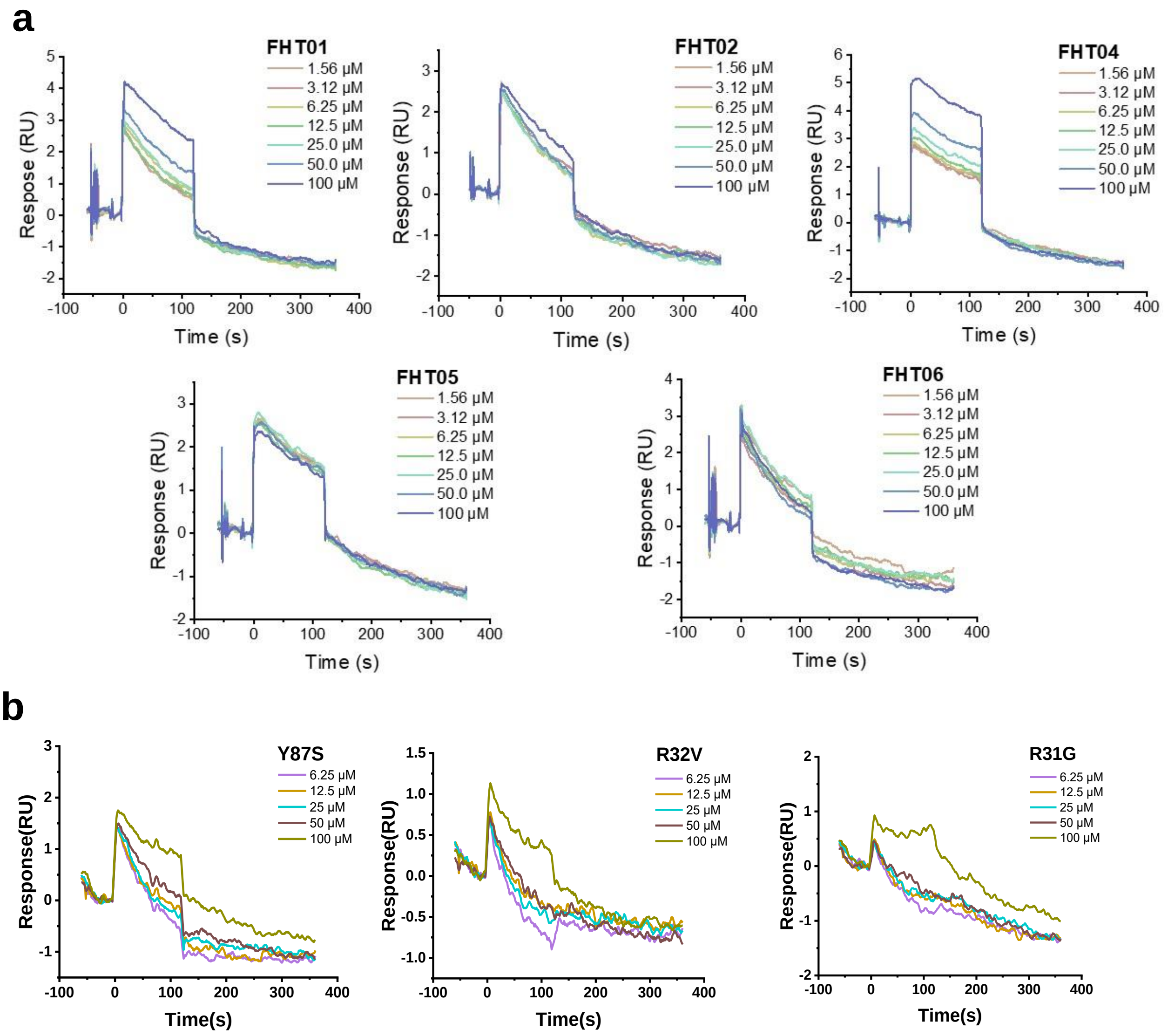


Figure S12: a.SPR direct binding test designed for cyclic peptide FHT03 and TNF$\alpha$ mutants which did not show significant responses, b.SPR direct binding test designed for FHT01, FHT02, FHT04, FHT05, FHT06 did not show significant responses.
